# Mechanism of Tethered Agonist-Mediated Signaling by Polycystin-1

**DOI:** 10.1101/2021.08.05.455255

**Authors:** Shristi Pawnikar, Brenda S. Magenheimer, Ericka Nevarez Munoz, Robin L. Maser, Yinglong Miao

## Abstract

Polycystin-1 (PC1) is an important unusual G-protein-coupled receptor (GPCR) with 11 transmembrane (TM) domains and its mutations account for 85% of cases of autosomal dominant polycystic kidney disease (ADPKD). PC1 shares multiple characteristics with Adhesion GPCRs. These include a GPCR proteolysis site that autocatalytically divides these proteins into extracellular, N-terminal and membrane-embedded, C-terminal (CTF) fragments, and a tethered agonist (TA) within the N-terminal stalk of the CTF that is suggested to activate signaling. However, the mechanism by which a TA can activate PC1 is not known. Here, we have combined functional cellular signaling experiments of PC1 CTF expression constructs encoding wild type, stalkless and three different ADPKD stalk variants with all-atom Gaussian accelerated molecular dynamics (GaMD) simulations to investigate TA-mediated signaling activation. Correlations of residue motions and free-energy profiles calculated from the GaMD simulations correlated with the differential signaling abilities of wild type and stalk variants of PC1 CTF. They suggested an allosteric mechanism involving residue interactions connecting the stalk, Tetragonal Opening for Polycystins (TOP) domain and putative pore loop in TA-mediated activation of PC1 CTF. Key interacting residues such as N3074-S3585 and R3848-E4078 predicted from the GaMD simulations were validated by new mutagenesis experiments. Together, these complementary analyses have provided novel insights into a TA-mediated activation mechanism of PC1 CTF signaling, which will be important for future rational drug design targeting PC1.

**Significance Statement:** Mutations of polycystin-1 (PC1) are the major cause (85% of cases) of autosomal dominant polycystic kidney disease (ADPKD), which is the fourth leading cause of kidney failure. PC1 is thought to function as an atypical GPCR, yet the mechanism by which PC1 regulates G-protein signaling remains poorly understood. A significant portion of ADPKD mutations of PC1 encode a protein with defects in maturation or reduced function that may be amenable to functional rescue. In this work, we have combined complementary biochemical and cellular assay experiments and accelerated molecular simulations, which revealed a novel allosteric transduction pathway in activation of the PC1 CTF. Our findings shall facilitate future rational drug design efforts targeting the PC1 signaling function.

## Introduction

Autosomal dominant polycystic kidney disease (ADPKD) is caused by mutations of the *PKD1* (85% of cases) or *PKD2* (15%) gene (1), which encode the proteins, polycystin-1 (PC1) and polycystin-2 (PC2), respectively. Together, PC1 and PC2 comprise a membrane receptor-ion channel signaling complex, involved in mechanosensing, signaling and intracellular [Ca^2+^] regulation (2-4). The PC1-PC2 complex is required for maintaining cellular homeostasis and differentiation (5, 6). Recent advancements in cryo-EM and structural studies of membrane proteins have led to the first glimpse of the PC1-PC2 complex (PDB: 6A70) (7), a heterotetramer consisting of 1 PC1 and 3 PC2 protein subunits in which the last 2 transmembrane (TM) domains of each protein form the ion channel pore (8), surrounded by the TOP (Tetragonal Opening of Polycystins) domains from the largest extracellular loop of each subunit.

PC1 is a large protein composed of 11 transmembrane (TM) domains and a short C-terminal cytosolic tail (C-tail) (9, 10). Its N-terminal, extracellular region consists of multiple binding domains that mediate cell-cell and cell-matrix interactions, Wnt binding and PC1-PC2 channel activation (9, 11-15). The C-tail initiates a number of signaling cascades (1), including the binding and activation of heterotrimeric G proteins, suggesting that PC1 functions an atypical type of G protein-coupled receptor (GPCR) (16-23). Recently, the regulation of G protein signaling by PC1 was shown to be critical for the prevention of renal cyst formation in mice (24, 25). However, we know very little about the molecular mechanism involved.

PC1 and the Adhesion family of GPCRs (ADGRs) share a conserved G protein-coupled receptor proteolysis site (GPS) motif located just proximal to their first TM domain that undergoes an intramolecular, self-catalyzed cleavage resulting in an N-terminal, extracellular fragment (**NTF**) and a C-terminal, membrane-embedded fragment (**CTF**) that remain non-covalently attached (26-31). GPS cleavage is known to be important for PC1 function, maturation and trafficking (27, 30, 32-35), and essential for preventing renal cystogenesis (32, 36). For the ADGRs, GPS cleavage and the association-dissociation of the NTF and CTF subunits are thought to be involved in regulation of G protein signaling (37-40). In the tethered agonist (TA) model of ADGRs, dissociation of the NTF results in exposure of a short, extracellular ‘stalk’ at the N-terminus of the CTF subunit which binds the extracellular loops and 7-TM bundle causing conformational changes that activate G protein signaling (41-44). We recently demonstrated that PC1 utilizes an ADGR-like TA for constitutive activation of an NFAT promoter-luciferase reporter (45). This work showed that the CTF subunit alone has greater signaling ability than full-length PC1, is dependent on the presence of the stalk and is affected by ADPKD-associated missense mutations within the stalk region. Furthermore, synthetic peptides derived from the PC1 stalk sequence could stimulate a stalkless CTF mutant to activate the NFAT reporter.

In the current study, we have combined complementary biochemical and cellular functional experiments and Gaussian accelerated molecular dynamics (GaMD) simulations to understand the dynamic structure-function relationships underlying the TA-mediated signaling of PC1 CTF. GaMD is an enhanced sampling computational technique that works by adding a harmonic boost potential to smooth the potential energy surface (46). GaMD greatly reduces energy barriers and accelerates biomolecular simulations by orders of magnitude. GaMD does not require pre-defined collective variables or reaction coordinates. Compared with the enhanced sampling methods that rely on careful selection of the collective variables, GaMD is of particular advantage for studying complex biological processes such as activation of the PC1 CTF. Moreover, because the boost potential exhibits a near-Gaussian distribution, free energy profiles can be properly recovered through cumulant expansion to the second order (46). Applications of GaMD have revealed mechanisms of protein folding and conformational changes, ligand binding, protein-protein/membrane/nucleic acid interactions and carbohydrate dynamics (47).

Here, a computational model of PC1 CTF was generated by extracting the protein from cryo-EM structure of the PC1-PC2 complex (PDB: 6A70) (7). Homology modeling was applied using I-TASSER (48) to add the missing regions in PC1 CTF, including the stalk, pore loop (PL) and C-tail (**Figs. 1A** and **S1**; see details in **SI Methods**). Parallel *in vitro* cellular signaling experiments and GaMD simulations were carried out on the wild type (WT), stalkless and ADPKD-associated stalk mutants of the PC1 CTF. Most of the PC1 CTF stalk variants exhibited reduced ability in activation of the intracellular NFAT reporter compared with the WT CTF. All-atom GaMD simulations revealed highly correlated residue motions and interactions between the stalk-TOP-PL domains, which appeared to be important for activation of the PC1 CTF. A number of simulation-predicted residue interactions in the PC1 CTF were further investigated by mutagenesis and cellular signaling experiments. The combined experiments and simulations have thus provided the first mechanistic insights into TA-mediated signaling by the PC1 CTF.

**Figure 1.**
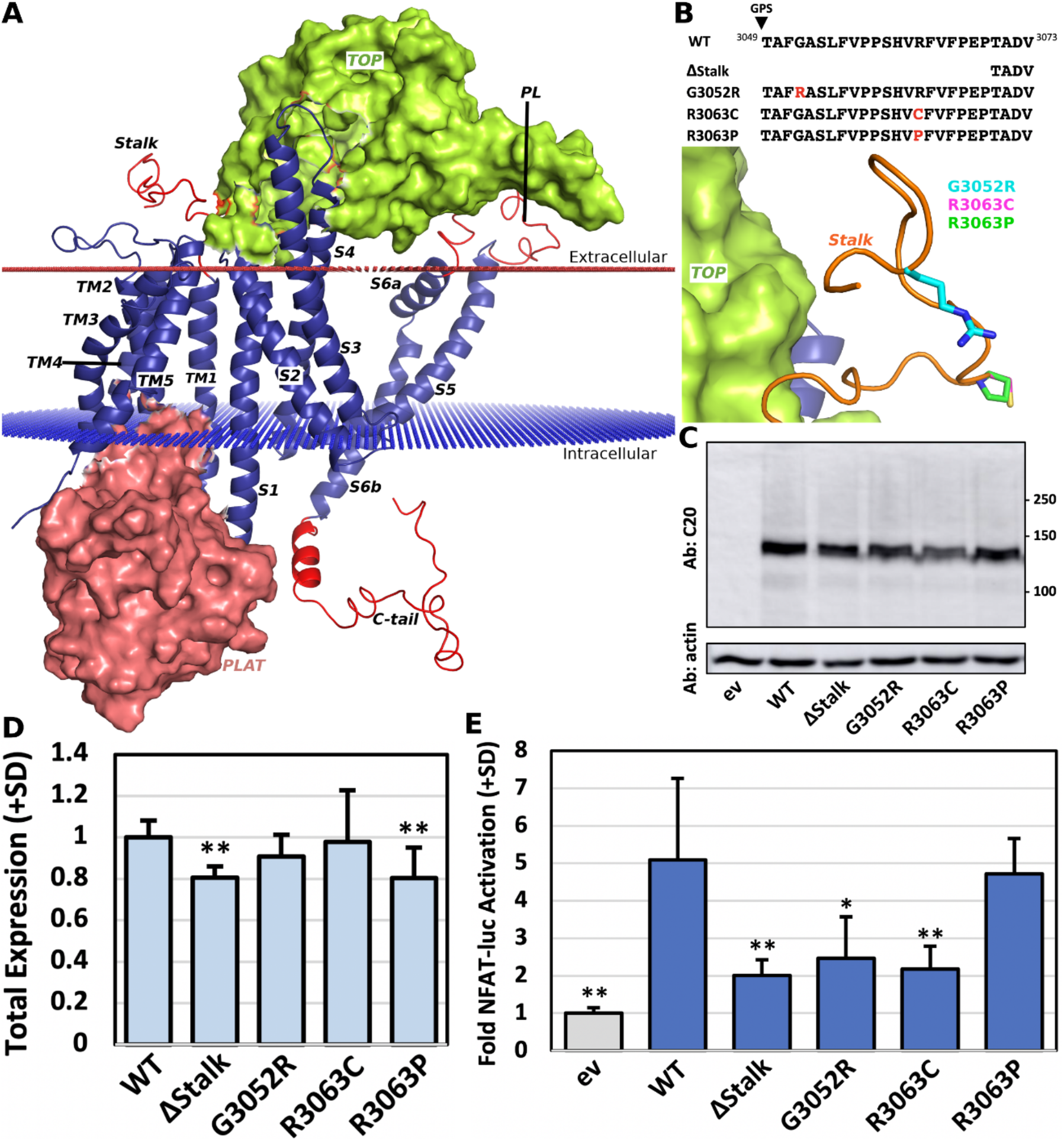
Atomic structure of PC1 CTF and experimental effects of stalk region variants on signaling to NFAT reporter. (A) Atomic structure of the PC1 CTF as extracted from the cryo-EM structure of PC1-PC2 complex (PDB: 6A70). The missing regions added through homology modeling, including Stalk, Pore Loop (PL) and C-tail, are highlighted in red. The TOP and PLAT domains are shown in green and pink surfaces, respectively. (B) Sequence alignment of the stalk region in wild type (WT), stalkless (ΔStalk), and the G3052R, R3063C and R3063P ADPKD missense mutants of human PC1 CTF and structural view of PC1 stalk variants with the mutated residues highlighted as sticks. The sequence alignment shows residue numbers (superscripted), GPS cleavage site (arrow), and ADPKD-associated missense mutations/polymorphisms (red). (C) Representative western blot of WT and mutant CTF proteins from one of the experiments summarized in (D). Blot was originally probed with C20 antibody and re-probed with antibody against actin. (D) Relative, average total expression level (+SD) for WT and mutant CTF constructs from signaling transfections represented in (E). (E) Average fold NFAT reporter activation (+SD) relative to empty vector (ev) for WT and mutant CTF proteins from 3-5 separate transfection experiments with n=3 wells/expression construct/experiment. *, p<0.005; **, p≤0.0001 relative to WT CTF levels.

## Results

### Stalk variants of the PC1 CTF exhibited reduced signaling activity in cellular assays

We first compared the ability of different expression constructs of human PC1 CTF to activate signaling to an NFAT reporter in transiently transfected HEK293T cells (**Fig. 1**). CTF expression constructs included the WT, stalkless (ΔStalk, with deletion of the first 21 N-terminal residues of CTF) and 3 missense mutants within the stalk region (G3052R, R3063C, R3063P) that are associated with ADPKD (https://pkdb.mayo.edu) (**Fig. 1B**). Semi-quantitative Western blot analyses showed that the deletion (ΔStalk) and ADPKD missense mutations of the stalk region had minimal effect (≤ 20%) on expression levels of the PC1 CTF (**Fig. 1C-1D**). However, the mutants exhibited distinct levels of activity in the NFAT reporter assay (**Fig. 1E**). WT CTF was able to activate the NFAT reporter by 5.09-fold relative to the empty expression vector (ev) control. In comparison, the ΔStalk, G3052R and R3063C variants of CTF activated the NFAT reporter only by 2.01-, 2.46- and 2.18-fold relative to ev control, respectively. These three stalk variations thus significantly reduced activity of the PC1 CTF in the NFAT reporter activation by ∼50-60% compared with the WT. Finally, the R3063P mutation resulted in only a minor reduction in NFAT reporter activation (4.71-fold relative to ev) compared to the WT CTF, which was not statistically significant (P = 0.577). We hypothesized that reductions observed in NFAT reporter activation of the ΔStalk, G3052R and R3063C variants of CTF were likely due to alterations in structural dynamics of the protein.

### GaMD simulations revealed reduced correlations in residue motions and disrupted domain interactions in the stalk variants of PC1 CTF compared with the wild type

In parallel with the mutagenesis and cellular assay experiments, we performed all-atom GaMD simulations on the WT and the ΔStalk, G3052R and R3063C variants of PC1 CTF (**SI Methods**). Three independent 1000 ns GaMD production trajectories were obtained on each system. The correlation matrix of residue motions was calculated from each trajectory and averaged over the three GaMD simulations for each system (**Fig. 2**). The corresponding standard errors of differences between the mutant and WT CTF systems were mostly smaller than 0.25 (**Fig. S2**). Important regions with significant difference in residue correlations between the mutant and WT CTF systems were circled in **Fig. 2**. In the WT CTF, high correlations were observed between the Stalk-TOP and TOP-PL domains (**Fig. 2A**). Notably, residues 3049-3074 in Stalk and 3699-3732 in TOP exhibited correlations > 0.5, and similarly for residues 3714-3880 in TOP and 4051-4080 in PL of the WT CTF. In comparison, correlations between residue motions in the stalk-TOP and TOP-PL domains were significantly reduced in the ΔStalk (**Fig. 2B**), G3052R (**Fig. 2C**) and R3063C (**Fig. 2D**) systems of the PC1 CTF compared with the WT (**Fig. 2A**). Residue correlations in the Stalk and TOP domains of the stalk variants of PC1 CTF showed values in the range of 0-0.25, being significantly lower than those in the WT (i.e., 0.25-0.75). Similarly, residue correlations in the TOP and PL domains of the ΔStalk and the ADPKD mutant systems also showed lower values of 0-0.35, as compared with those of the WT (0.25-0.75). The loss of correlated residue motions in the Stalk-TOP and TOP-PL domains observed in the stalk variants during GaMD simulations suggested that these domains were likely crucial for TA-mediated signaling in PC1 CTF, and therefore, were further investigated.

**Figure 2.**
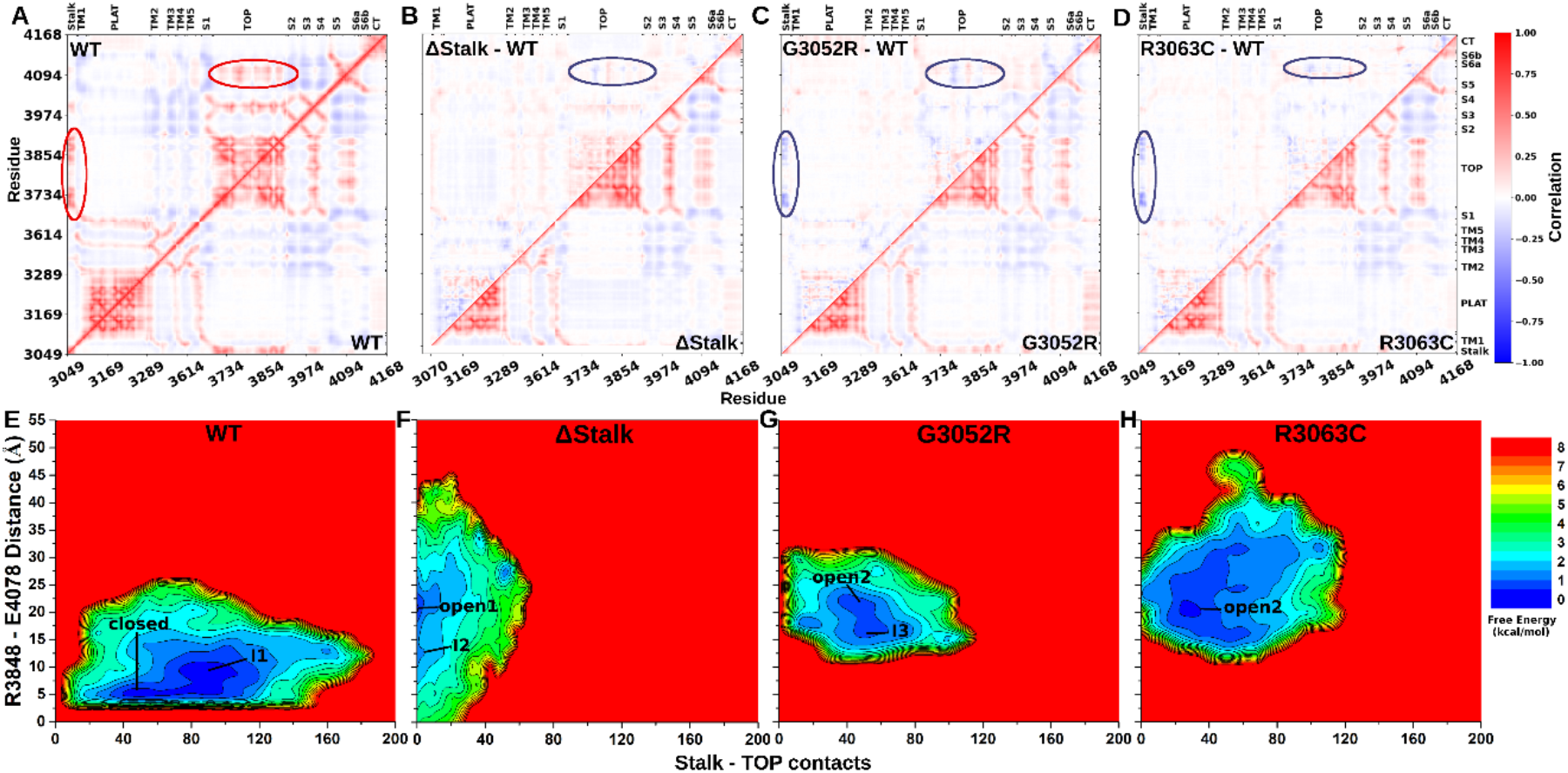
Reduced correlations in residue motions and disrupted domain interactions in the stalk variants of PC1 CTF compared with the wild type. (A) The correlation matrix of residue motions averaged over three GaMD simulations of the WT PC1 CTF. High correlations between Stalk-TOP and TOP-PL domains are highlighted in red circles. (**B-D**) The average correlation matrix of residue motions (lower triangle) and corresponding differences relative to the WT (upper triangle) calculated from three GaMD simulations of the (**B**) ΔStalk, (**C**) G3052R and (**D**) R3063C systems of PC1 CTF. Important regions with statistically significant differences (p<0.05) are highlighted in blue circles. (**E-H**) 2D free energy profiles of the (**E**) WT, (**F**) ΔStalk, (**G**) G3052R and (**H**) R3063C systems of PC1 CTF regarding the number of atom contacts between the Stalk and TOP domains and the R3848-E4078 distance (the CZ atom in R3848 and the CD atom in E4078) calculated from the GaMD simulations. Residue contacts were calculated for heavy atoms with a distance cutoff of 4 Å. Important low-energy conformational states are identified, including the “Closed”, “Intermediate I1”, “Intermediate I2”, “Intermediate I3”, “Open1” and “Open2”.

During the GaMD simulations, a salt bridge was frequently formed between residues R3848-E4078 in the WT CTF, but rarely observed in the ΔStalk and completely absent in the ADPKD mutant systems as shown in **Fig. S3**. Therefore, the R3848-E4078 ionic interaction appeared to play an important role in facilitating stalk-mediated signaling of the WT CTF. In addition, a hydrogen bond was formed between residue N3074 at the base of the stalk region and S3585 near the extracellular end of TM5 in two of the three GaMD simulations of WT CTF, but was intermittently observed in the stalk variant mutants (**Fig. S4**).

Next, we calculated 2D free energy profiles of the WT and stalk variants of PC1 CTF by reweighting of the GaMD simulations (**Fig. 2E-2H**). The R3848-E4078 residue distance and the number of contacts between the Stalk and TOP domains were selected as the reaction coordinates. Two low-energy conformational states were identified from the free energy profile of the WT CTF and designated as “Closed” and “Intermediate I1” (**Fig. 2E**). In the WT CTF, the stalk region was found to interact considerably with the TOP domain, for which their number of atom contacts ranged ∼40-120 (**Figs. 2E** and **S5A**). In the “Closed” state, a salt bridge was formed between R3848-E4078 with a distance of ∼3.9 Å between the residue charge centers (the CZ atom in R3848 and the CD atom in E4078). The “Intermediate I1” state showed an increased number of ∼60-120 contacts between Stalk-TOP domains, however, the R3848-E4078 salt bridge became broken at ∼8 Å distance between the residue charge centers (**Fig. 2E**). In addition to the CHARMM36m force field, we performed three independent 500 ns GaMD simulations on the WT PC1 CTF system using the AMBER FF19SB force field. Similarly, high residue correlations were observed between the Stalk-TOP and TOP-PL domains (**Fig. S6A**). Low-energy conformational states including “closed” and “I1” were also identified from the 2D free energy profile (**Fig. S6B**) regarding the Stalk-TOP contacts and the R3848-E4078 distance (**Fig. S6C**).Overall, consistent results were obtained from GaMD simulations using different force fields on the WT CTF.

In the ΔStalk system, two low-energy conformational states (“Intermediate I2” and “Open1”) were identified from the free energy profile (**Fig. 2F**). “Intermediate I3” low-energy state was identified for the G3052R mutant along with “Open2” (**Fig. 2G**), which was also sampled in the R3063C system (**Fig. 2H**). The R3848-E4078 salt bridge became broken in the I2, I3, Open1 and Open2 low-energy states (**Fig. 2F-2H**) and the number of contacts between the Stalk-TOP domains centered around 0, ∼50 and ∼30 in the ΔStalk (**Fig. 2F**), G3052R (**Fig. 2G**) and R3063C (**Fig. 2H**) systems, respectively.

Additionally, we calculated free energy profiles of individual GaMD simulations for each system, including the WT, ΔStalk, G3052R, R3063C and R3063P (**Figs. S7 to S11)**. For the WT CTF, both Sim1 and Sim2 sampled the “closed” low-energy state (**Fig. S7A-B**) and the “I1” state was observed mainly in Sim3 (**Fig. S7C)**. For the ΔStalk CTF, both Sim2 and Sim3 sampled the “open1” state and the “I2” state was observed in Sim1 and Sim3 (**Fig. S8**). For the G3052R mutant, both Sim2 and Sim3 sampled the “open2” state and Sim1 sampled the “I3” state (**Fig. S9**). The R3063C mutant sampled the “open2” state in all three GaMD simulations (**Fig. S10**). Finally, the R3063P mutant sampled the intermediate “I1” (**Fig. S11A**) and “I3” (**Fig. S11B-C**) states. Despite the variations, the free energy profiles of individual GaMD simulations showed consistent results as compared with those calculated with all the simulations combined for each system. The R3848-E4078 salt bridge tended to be broken between the TOP-PL domains and the Stalk-TOP contacts decreased in the stalk variants.

### Distinct low-energy conformations of the wild type and stalk variants of PC1 CTF

In the “Closed” low-energy conformation identified for the WT CTF, a number of polar interactions were identified between the Stalk and TOP domains, including a hydrogen bond between N3074-S3585 (**Figs. 3B** and **S4A**), T3049-E3708 (**Figs. 3A** and **S12A**) and a salt bridge between E3068-R3700 (**Figs. 3A** and **S13A**). The distance between the OG1 atom of T3049 and the CD atom of E3708 was calculated as ∼3.6 Å, and ∼4.7 Å distance between the charge centers of E3068 (the CD atom) and R3700 (the CZ atom) (**Fig. 3A**). Note that the T3049-E3078 hydrogen bonding interaction was present in only the WT CTF (**Fig. S12**), while the E3068-R3700 salt bridge was formed in both the WT and R3063P mutant CTF (**Fig. S13**). The N3074-S3585 hydrogen bond was observed to form very stably in the WT CTF system, however, was intermittently present in the CTF mutants. The free energy minimum distance between the CG atom in N3074 and the OG atom in S3585 was found to be ∼3.9 Å (**Fig. 3B**). Furthermore, a salt bridge was formed between R3848-E4078 from TOP and PL domains with a distance of ∼3.9 Å between the residue charge centers (**Figs. 3C** and **S3A**).

**Figure 3.**
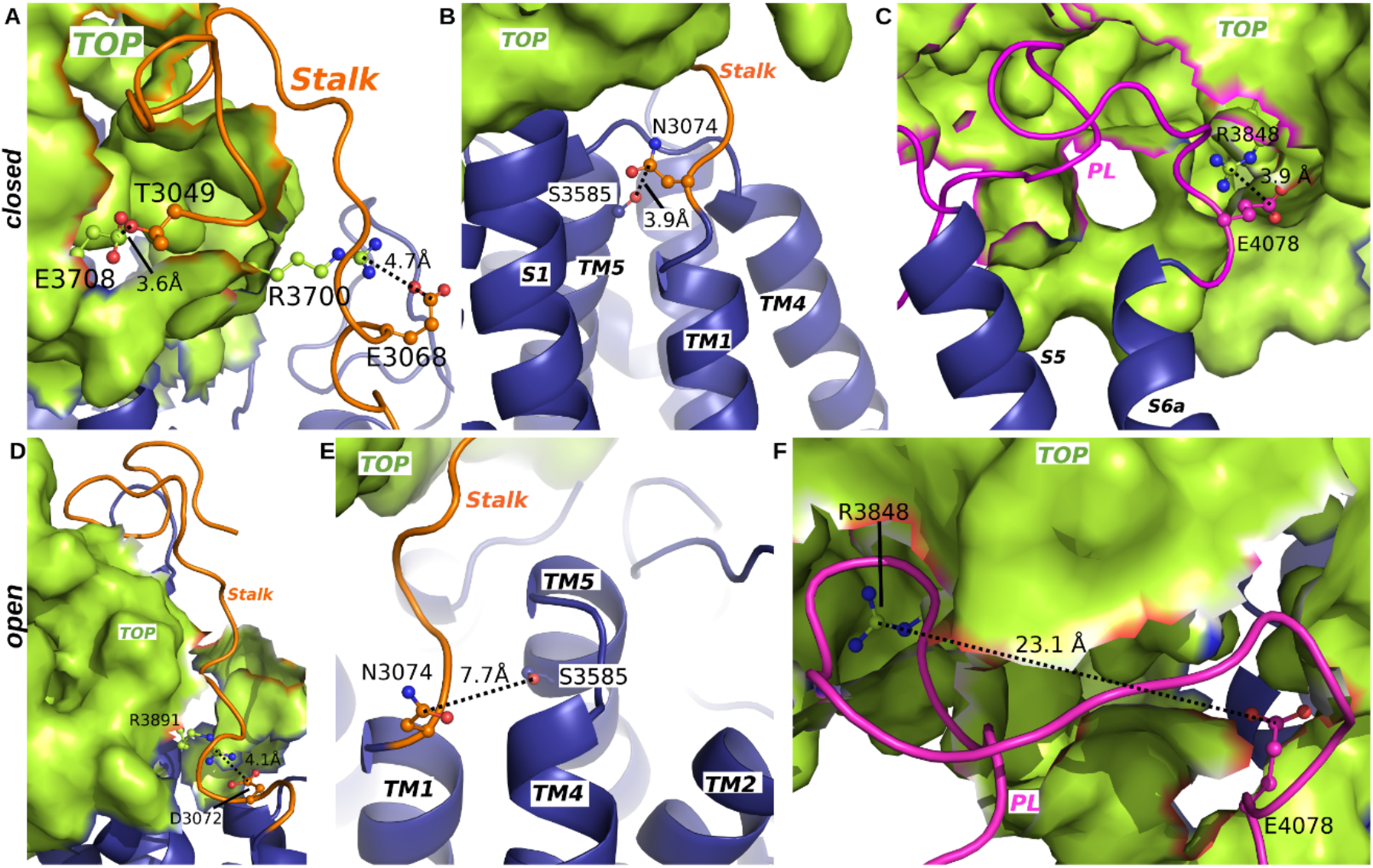
Distinct low-energy conformations of the WT and stalk variants of PC1 CTF. **(A-C)** Residue interactions observed in “Closed” protein conformation between (A) Stalk-TOP, (**B**) Stalk-TM5 and (**C**) TOP-PL domains. (**D-F**) Residue interactions observed in “Open” conformation of the Stalk variants between (**D**) Stalk-TOP, (**E**) Stalk-TM5 and (**F**) TOP-PL domains. The Stalk (orange cartoon), transmembrane helices (blue cartoon), TOP domain (green surface) and pore loop (magenta cartoon) are labeled in the PC1 CTF. Residue interactions are represented in ball and stick.

In the “Open1” and “Open2” low-energy conformational states identified from free energy profiles of the ΔStalk, G3052R and R3063C PC1 CTF, residue D3072 from the C-terminal end of the Stalk region formed a salt bridge with R3891 in the TOP domain (**Figs. 3D** and **S14**). The D3072-R3891 salt bridge interaction was formed for more extended periods of time during GaMD simulations of these three systems as compared to the WT CTF and R3063P mutant. However, the N3074-S3585 hydrogen bond became broken with ∼7.7 Å distance between the residue charge centers (CG atom in N3074 and OG atom in S3585) (**Fig. 3E**). The salt bridge interaction connecting residues R3848-E4078 from TOP and PL domains also became broken with a large distance of 23.1 Å between the residue charge centers (**Fig. 3F**), although fluctuations were observed during the GaMD simulations (**Fig. S3B-D**).

### Intermediate conformational states of PC1 CTF observed from GaMD simulations of the ΔStalk, G3052R and R3063P mutant

In addition, three independent 1000 ns GaMD simulations were performed on the R3063P ADPKD missense mutant, which showed only a minor reduction in NFAT reporter activation compared to WT PC1 CTF (**Fig. 1E**). Correlations between residues motions in the stalk-TOP and TOP-PL domains were slightly reduced in the R3063P mutant compared with the WT (**Fig. 4A**). Residue motions in the Stalk and TOP domains of the R3063P PC1 CTF system showed correlation values ranging between 0.25-0.5 as compared to correlation values of 0.25-0.75 for the WT system. Similarly, residue motions in the TOP and PL domains of the R3063P PC1 CTF system ranged between 0.25-0.5, which were slightly lower than those of the WT (0.25-0.75).

**Figure 4.**
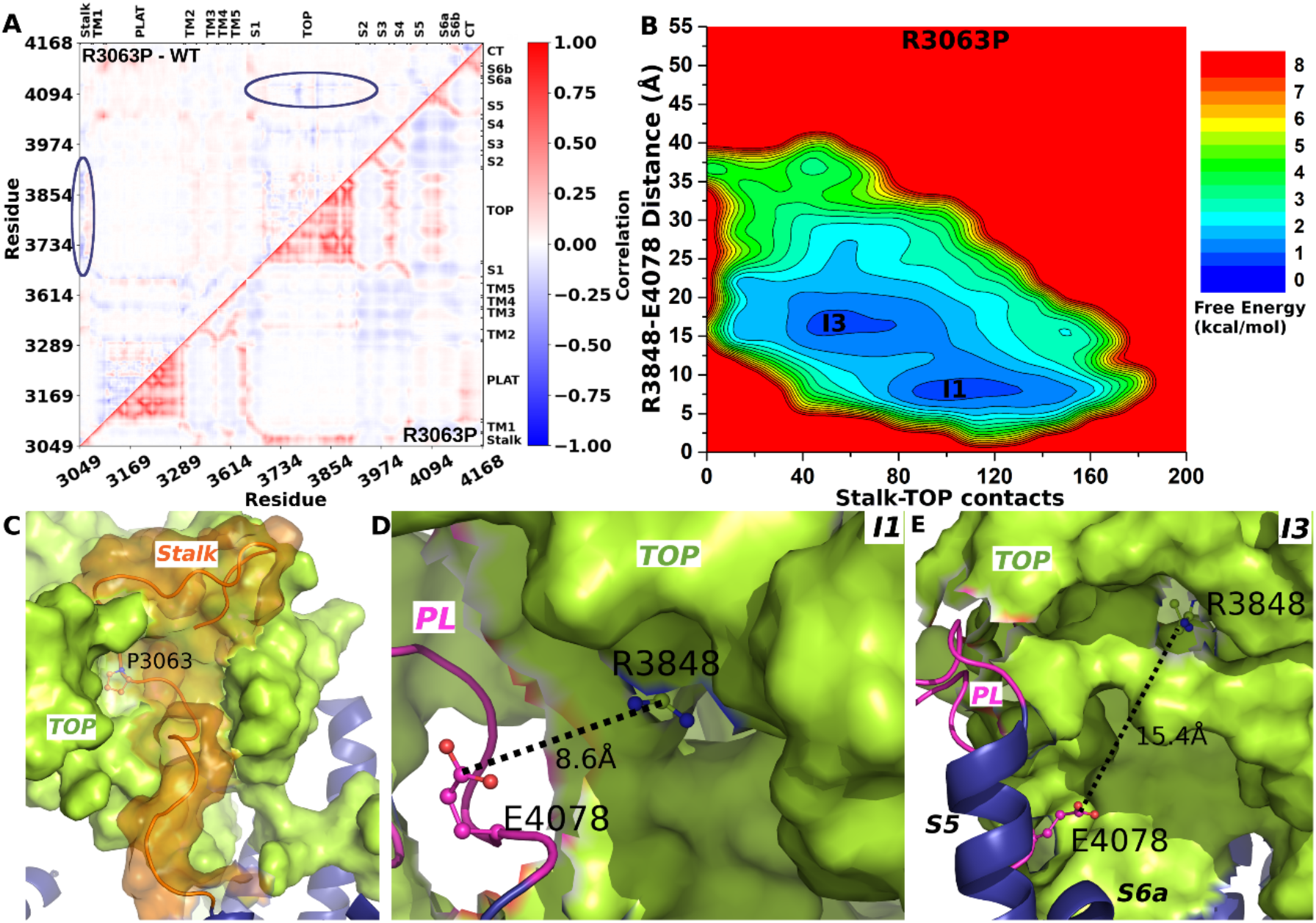
Intermediate conformational states of PC1 CTF observed from GaMD simulations of the R3063P mutant. (**A**) The average correlation matrix of residue motions (lower triangle) and corresponding differences relative to the WT (upper triangle) calculated from three GaMD simulations of the R3063P mutant. (**B**) 2D free energy profile of the R3063P mutant regarding the number of contacts between the Stalk and TOP domains and the R3848-E4078 distance calculated from the GaMD simulations. Important low-energy conformational states are identified, including the “Intermediate I1” and “Intermediate I3”. (**C**) Structural conformation of Stalk-TOP interaction for the R3063P mutant. (D) Residue interactions between TOP-PL as observed in the “Intermediate I1” conformational state. (E) Residue interactions between TOP-PL as observed in the “Intermediate I3” conformational state.

A 2D free energy profile was further calculated for the R3063P mutant of PC1 CTF regarding the number of contacts between Stalk-TOP and R3848-E4078 residue distance (**Fig. 4B**). Two low-energy conformational states were identified from the free energy profile, i.e., “Intermediate I1” and “Intermediate I3”. Note that the “I1” and “I3” states were observed in the WT (**Fig. 2E**) and G3052R mutant (**Fig. 4B**) of PC1 CTF, respectively. Overall, the R3848-E4078 salt bridge became broken in these two intermediate conformational states, but the residue distance was shorter than in the “Open1” and “Open2” states observed with the ΔStalk, G3052R and R3063C mutant systems (**Fig. 2F-H**). Additionally, an “Intermediate I2” low-energy conformational state was observed in the ΔStalk system. It showed zero contacts between the Stalk-TOP due to absence of the Stalk and the R3848-E4078 salt bridge became broken at ∼13-15 Å distance (**Fig. 2F**). In comparison, the “Intermediate I1” state showed increased number of contacts between Stalk-TOP ranging ∼90-130 (**Fig. 4B-4C**) and the R3848-E4078 distance decreased to 7-9 Å (**Fig. 4B** and **4D**). The “Intermediate I3” state showed ∼40-80 contacts between Stalk-TOP (**Fig. 4B**) and the R3848-E4078 distance increased to ∼15.4 Å (**Fig. 4B** and **4E**).

### Experimental validation of simulation-predicted residue interactions that are important for WT stalk-mediated activation of PC1 CTF

To determine whether newly discovered residue interactions in the GaMD simulations, i.e., the N3074-S3585 hydrogen bond and R3848-E4078 salt bridge, are important for signaling of the WT CTF, these residues were mutated, one at a time, in the CTF expression construct and assessed for signaling to the NFAT reporter (**Fig. 5A**). Activation of the NFAT reporter was significantly reduced from 6.10-fold for WT CTF to 2.00-fold (P < 0.0001) for N3074A and 3.69-fold (P = 0.0066) for S3585A CTF mutants. Likewise, the ability of activating the NFAT reporter was significantly reduced for the R3848E mutant (2.15-fold, P =0.0001) with respect to WT CTF, although reporter activation was only slightly reduced for the E4078R mutant (5.57-fold, P = 0.6021). To determine if changes in NFAT reporter activation by any of the mutants was due to loss of expression or inability to traffic, both the total (**Fig. 5B-5C**) and cell surface (**Fig. 5D-5E**) expression levels were investigated. Semi-quantitative Western blot analyses show total expression levels of the CTF mutants were reduced to 0.79 for N3074A, 0.71 for S3585A and R3848E, and 0.51 for E4078R of WT CTF level, while cell surface expression levels were increased for CTF-N3074A and -S3585A (1.20- and 1.41-fold, respectively) and decreased for R3848E and E4078R (0.51 and 0.71, respectively) relative to WT CTF.

**Figure 5.**
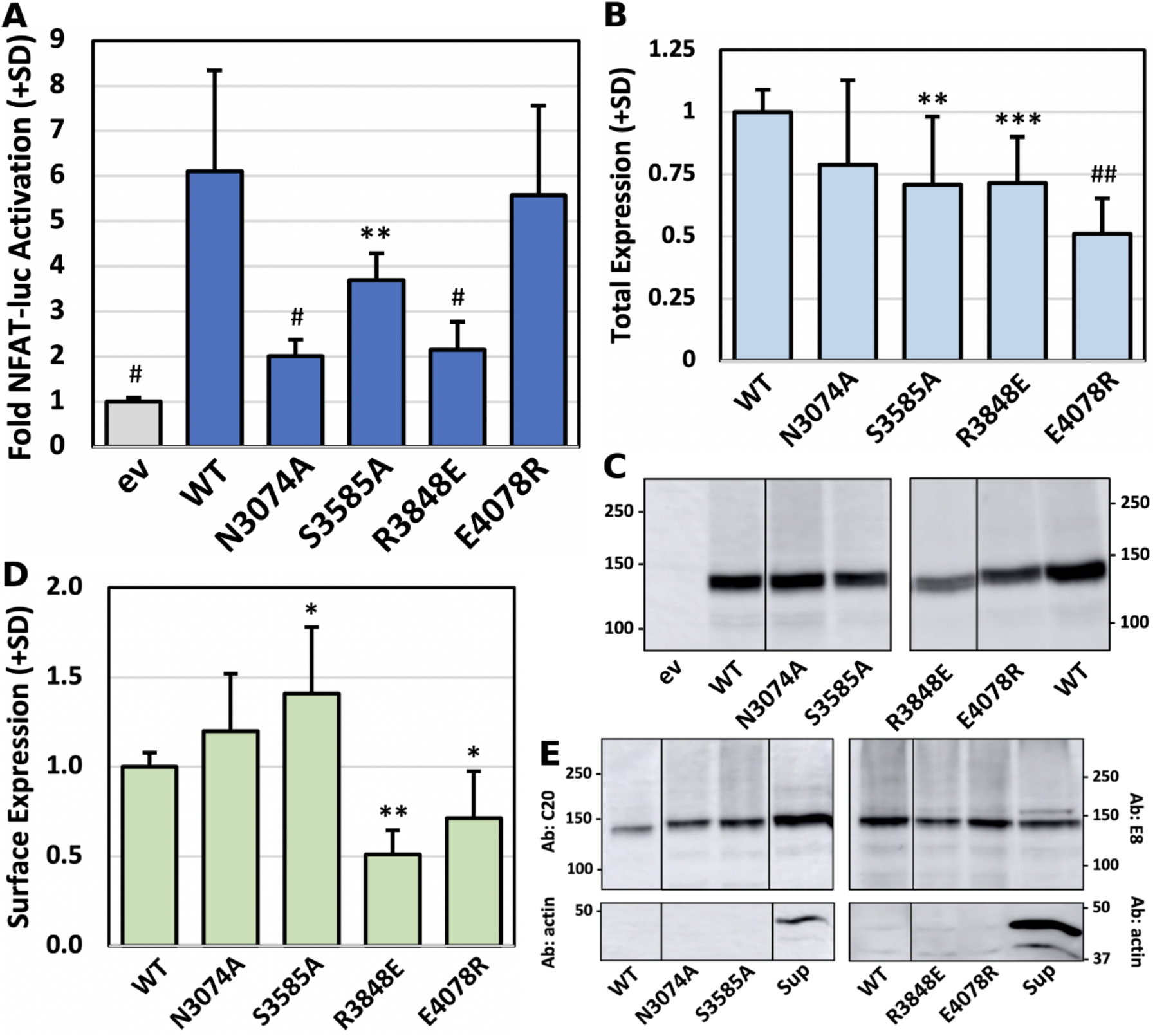
Experimental analyses of GaMD simulation-predicted residue interactions between TOP-PL domains important for WT stalk-mediated activation of PC1 CTF. (A) Average fold NFAT reporter activation (+SD) relative to empty vector (ev) for WT and GaMD mutant CTF proteins from 3 separate transfection experiments with n=3 wells/expression construct/experiment. (B) Average relative total expression level (+SD) for WT and GaMD mutant CTF proteins from experiments in (A). (C) Representative Western blots of WT and GaMD mutant CTF proteins from one of the experiments represented in (B) probed with C20 antibody. Images are from two separate Western blots. Removal of intervening lanes is indicated by solid line. Blots were originally probed with C20 antibody and reprobed for actin. (D) Average relative cell surface expression level (+SD) for WT and GaMD mutant CTF proteins from 2-6 separate surface biotinylation experiments. (E) Representative Western blots of the surface biotinylation analyses of WT and GaMD mutant proteins from one of the experiments represented in (D). Supernatant (Sup) lane is from the pulldown with WT CTF. Images are from two separate blots. Removal of intervening lanes indicated by solid lines. Blots were originally probed with C20 or E8 antibody as indicated and reprobed for actin. *, p<0.05; **, p≤0.01; *** p<0.005; #, p<0.0005; and ##, p<0.0001 relative to WT CTF levels.

To determine if the effects on PC1 CTF signaling and expression levels by the above residue mutations were random or specific to the proposed mechanism, we attempted to made additional CTF expression constructs with residue substitutions designed to not disrupt our predicted residue interactions or to not interfere with the proposed allosteric mechanism, i.e., neutral mutations. These putative neutral mutations consisted of a conservative substitution of residue N3074 (N3074Q), replacement of a charged residue within the TOP domain located away from proposed interaction sites with the stalk or the PL (R3835A), and a reportedly benign substitution within the PC1 CTF stalk (F3066L) (https://pkdb.mayo.edu/variants). The signaling ability and expression levels of neutral CTF mutants were analyzed and compared to WT CTF (**Fig. S15**). All three mutants had near WT total and surface expression levels (**Fig. S15B, C**). N3074Q greatly inhibited signaling to the NFAT reporter, while R3835A had a much smaller effect and F3066L had no significant effect on signaling to the NFAT reporter (**Fig. S15A**).

## Discussion

In this study, we have combined complementary site-directed mutagenesis and cellular experimental assays with all-atom GaMD simulations to decipher the TA-mediated activation mechanism of the PC1 CTF. The ΔStalk, G3052R and R3063C variants significantly reduced activation of the NFAT reporter compared with the WT CTF, unlike the R3063P mutant. Correlation matrices calculated from GaMD simulations of the WT and mutant CTF systems revealed alterations in residue motions between the stalk and TOP and the TOP and PL domains that were highly consistent with the results of the cellular signaling assays. Further analyses of the GaMD simulations revealed an allosteric transduction pathway connecting the stalk-TOP-PL domains, which was important for activation of the WT CTF in the “Closed” conformational state (**Fig. 6A**). This pathway and interactions between the stalk-TOP-PL domains were disrupted in the ΔStalk, G3052R and R3063C systems, leading to a distinct “Open” (Open1 and Open2) conformational state in the calculated free energy profiles (**Fig. 6B**). This was in contrast to the “Closed” and “Intermediate” (I1, I2 and I3) low-energy states of the WT, ΔStalk, G3052R and R3063P PC1 CTF.

**Figure 6.**
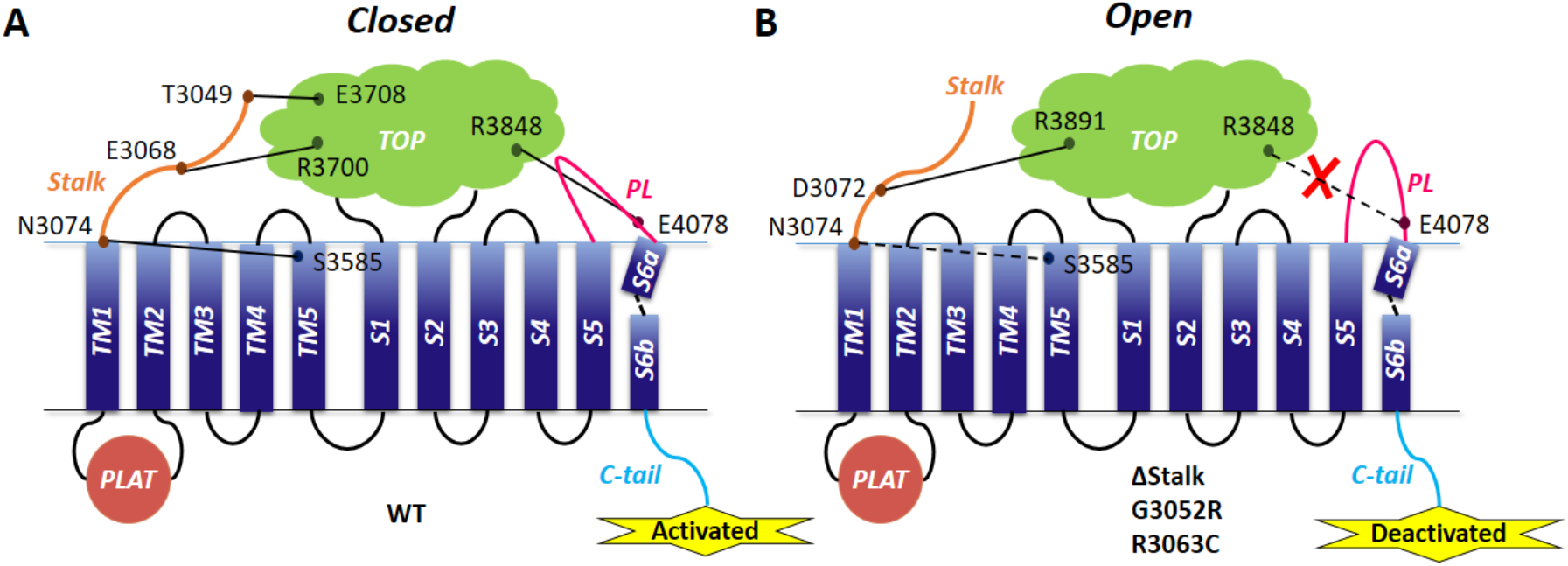
Model of the Stalk TA-mediated activation signaling of the PC1 CTF. (A) A “Closed” low-energy conformation is identified for the WT CTF, in which signal is transduced through an allosteric pathway connecting the Stalk-TOP-PL domains and ultimately to the C-tail for G protein activation. (B) Stalk variants (including ΔStalk, G3052R and R3063C) of CTF adopt a distinct “Open” conformation, which exhibits reduced residue interactions between the Stalk-TOP domains and broken salt bridge interaction of residues R3848 and E4078 between the TOP-PL domains, leading to significantly reduced signaling activity of these PC1 mutants.

It is important to note that the free energy profiles calculated from GaMD simulations of PC1 CTF were not fully converged since certain variations were observed among the individual simulations (**Figs. S7 – S11**). It is exceedingly difficult to calculate accurate free energy profiles for a large membrane protein such as PC1 CTF. Longer simulation lengths (e.g., 2000 ns) and a larger number of independent GaMD production simulations (e.g., ∼5-10) could potentially improve conformational sampling and free energy profiling calculations of the PC1 CTF. Nevertheless, similar low-energy conformational states could be identified from free energy profiles of current individual GaMD simulations as compared with those from all simulations combined for each system. Overall, mutations in the Stalk led to a reduced number of contacts between Stalk-TOP domains and breaking of the R3848-E4078 salt bridge between the TOP-PL domains in PC1 CTF. Newly found residue interactions such as N3074-S3585 and R3848-E4078, which were predicted to be important in the GaMD simulations, were supported by mutagenesis and cellular signaling experiments. It is important to note given the differences in complexity between the two experimental systems (i.e., cellular milieu versus defined *in silico* conditions) that these amino acid substitutions could alter unrelated or unintended structural or functional properties of the PC1 CTF (e.g., interactions with other proteins, delivery to specific plasma membrane domains, etc), and hence result in effects on signaling to the NFAT reporter. Nonetheless, our experiments and simulations combined have provided novel mechanistic insights into the TA-mediated activation of the PC1 CTF signaling activity.

GaMD simulations suggested a large number of contacts formed between the Stalk region and TOP domain in the WT CTF. Removal of the stalk region and single residue mutations G3052R and R3063C decreased the number of Stalk-TOP contacts and disrupted the R3848-E4078 salt bridge interaction between the TOP-PL domains. Notably, residues 3700-3730 exhibited high correlations (> 0.75) with residues 3820-3890 in the TOP domain, and similarly high correlations were evident between residues 3820-3850 and residues 3850-3890 (**Fig. S16**). Such high correlations of residue motions within the TOP domain likely contributed to the allosteric signal transmission of PC1 CTF from the Stalk-TOP interface to the TOP-PL interactions site involving the R3848-E4078 salt bridge (**Fig. S16B**). These observations suggest that the stalk interaction with the TOP domain could trigger signal activation being translated via the R3848-E4078 salt bridge interaction between the TOP-PL domains and ultimately to the C-tail of PC1, which contains the G protein activation motif (49).

GaMD simulations also revealed a hydrogen bond interaction between N3074 located at the base of the Stalk and S3585 in the extracellular domain of TM5 in WT CTF, which was intermittently present in the CTF mutants. The importance of this interaction in stalk-mediated signaling of PC1 CTF was experimentally investigated. Substitution of either residue with Ala to prevent their polar interaction resulted in a significant reduction in activation of the NFAT reporter. Together with the apparent weakening or loss of this interaction in the GaMD simulations of stalk variants, these results suggested that the N3074-S3585 hydrogen bond was likely important in anchoring the base of the Stalk for interactions with the TOP domain. Such a proposed role for the N3074-S3585 interaction may have been replaced by residue-residue contacts between D3072-R3891 that are more prominent in the ΔStalk, G3052R and R3063C mutants compared to WT CTF and the R3063P mutant (**Figs. 6** and **S14**). Furthermore, retention of the N3074-S3585 and E3068-R3700 interactions by the R3063P mutant appeared to be able to compensate for its stalk residue substitution and promote a high number of functional Stalk-TOP contacts, thus explaining its only slightly reduced ability in activating the NFAT reporter. Relevant to these findings, it was intriguing that both residues N3074 and E3068 were reported to be sites of ADPKD-associated mutations (PKD mutation database, https://pkdb.mayo.edu) and were recently shown to alter signaling to the NFAT reporter when mutated.

The importance of the R3848-E4078 salt bridge interaction between the TOP-PL domains was also experimentally tested through mutation-signaling experiments. The salt bridge was disrupted by mutating either residue to an oppositely charged residue (R3848E and E4078R), which was expected to cause repulsion between the TOP domain and PL. While the R3848E mutant showed significantly decreased activation of the NFAT reporter as compared to the WT, the signaling ability of the E4078R CTF mutant was only slightly decreased (**Fig. 5A**). Since the total and cell surface expression levels of CTF-E4078R were lower than WT CTF, one might argue that this mutant’s activity relative to its expression level was actually greater than WT CTF. In this regard, residue mutations that could even increase signaling capability of ADGRs have been identified (50, 51). A lack of correlation between surface expression levels and signaling activity of ADGRs has been also reported for GPR133/ADGRD1 (52). A possible explanation for the apparent lack of effect of the E4078R substitution in the PL on NFAT reporter activation could be an ability of the E4078R mutant to re-establish a fortuitous interaction with one of the many polar residues in the TOP domain, which maintained allosteric signaling of the Stalk-TOP interactions to the PL. On the other hand, the PL does not include any basic residues that would enable mutation of residue R3848 in the TOP domain to negatively charged Glu to interact, thus disrupting its interaction with the PL and resulting in the significantly reduced activity of the R3848E mutant. Regardless, these findings further support a model where the TOP-PL interactions are involved in stalk-/TA-mediated activation of signaling in PC1 CTF.

Attempts to introduce putative neutral mutations into CTF that would not disrupt the proposed allosteric TA-mediated activation mechanism in order to gauge the specificity of our mutation-signaling validation experiments yielded mixed results (**Fig. S15**). Substitution of N3074 with a similarly polar Gln residue dramatically reduced CTF signaling ability, replicating the results obtained with N3074A. As mentioned above, an ADPKD-associated mutation, N3074K, also inhibits CTF-mediated activation of the NFAT reporter (45), which may indicate that the size of the residue at position 3074 in addition to its polarity is strictly required and further underscores the importance of this residue in ‘setting up’ the ability of the stalk to efficiently interact with the TOP domain. R3835A, located at the top of the TOP domain in our model, had a small but significant effect on reporter activation. While we chose R3835 based on its location away from the stalk-TOP and TOP-PL interactions and its reduced conservation among PC1 of higher vertebrates (i.e., replaced by Gly in pig, dog, rat, mouse), the preponderance of polar and charged residues within the TOP domain could be important for transmission of the allosteric mechanism, as suggested by **Fig. S16**, and thus may have precluded a neutral effect by its substitution to Ala. Interestingly, F3066L, which is a common substitution in the PKD1 gene and predicted to be ‘likely benign’, did not inhibit CTF signaling to the NFAT reporter, thus supporting the specificity of amino acid substitutions for investigating the TA mechanism. Substitution of any residue within a protein, let alone a protein with an allosteric functional mechanism, has the potential to alter protein structure, and hence function, which emphasizes the need for additional molecular dynamics simulations to better understand the structure-function relationships of important proteins.

Shared features between PC1 and the ADGRs, including auto-proteolytic cleavage at the GPS motif and homology of the GAIN domain, led to the proposal and investigation of whether a TA is involved in PC1 CTF-mediated activation of an NFAT luciferase reporter (45). The current study suggests that the PC1 CTF utilizes a novel allosteric activation mechanism for its TA-mediated signaling activation. To date, neither the molecular mechanism of TA-mediated activation nor the binding pocket of the TA have been established for ADGRs, although a number of observations are supportive of an allosteric activation mechanism (53, 54). The involvement of extracellular loops and ends of TM domains in either binding of a tethered ligand or contributing to activation of signaling has been reported for a number of ADGRs including Gpr56/ADGRG1, Lphn1/ ADGRL1, GPR133/ADGRD1, and GPR64/ADGRG2 (51, 52, 55, 56). Recently, the binding pocket for the tethered ligand of protease-activated receptor 4 (PAR4) was also determined to involve residues in the TM3 and TM7 and ligand binding was regulated by extracellular loop 3 (57).

In summary, we have revealed novel Stalk-TOP and TOP-PL domain interactions, which played a significant role in the activation mechanism of the PC1 CTF. Our complementary mutagenesis and cellular signaling experiments and GaMD simulations on both the WT and signaling-deficient mutant CTF proteins have provided important insights into the mechanism of PC1 activation at an atomic level, which are expected to facilitate future rational drug design efforts targeting the PC1 signaling protein.

## Materials and Methods

We have performed transient transfections of HEK293T cells with WT and mutant CTF expression constructs of PC1 to assess their ability to activate signaling to an NFAT promoter luciferase reporter following a previously established protocol (58). PC1 CTF mutants, including the ΔStalk (a 21-residue deletion of the CTF stalk region), single residue substitutions (the ADPKD-associated mutations G3052R, R3063C, R3063P and F3066L), simulation suggested mutations N3074A, S3585A, R3848E, and E4078R, and potential neutral mutations (N3074Q, R3835A) were generated by site directed mutagenesis. To evaluate the potential effect of mutations in the PC1 CTF, total and cell surface expression levels were determined from total cell lysates and surface biotinylated proteins, respectively, from the transfected cells using SDS-PAGE and Western blot analysis. GaMD simulations were performed on the WT, ΔStalk, G3052R, R3063C and R3063P systems of the PC1 CTF, following a previously established protocol (59). The CHARMM36m parameter set (60) was used for the protein and lipids. NAMD2.12 (61) was used for initial energy minimization, thermalization and 20 ns conventional MD equilibration. The output files of cMD simulations using NAMD were converted to AMBER format using *ParmEd* tool AMBER package (62). GaMD simulations implemented in GPU version of AMBER 18 (46) were performed for the PC1 CTF and its mutant systems. The GaMD simulations involved a short conventional MD of 12 ns, 48 ns GaMD equilibration and finally three independent 1 μs GaMD production runs with randomized initial atomic velocities for each system. Details of mutagenesis, cellular signaling assays, surface biotinylation, Western blot analysis, GaMD simulations, system setup, and simulation analysis are provided in **SI**.

## Acknowledgments

This work used supercomputing resources with allocation award TG-MCB180049 through the Extreme Science and Engineering Discovery Environment (XSEDE), which is supported by National Science Foundation grant number ACI-1548562, and project M2874 through the National Energy Research Scientific Computing Center (NERSC), which is a U.S. Department of Energy Office of Science User Facility operated under Contract No. DE-AC02-05CH11231, and the Research Computing Cluster at the University of Kansas. This work was supported in part by National Institutes of Health (DK123590), Department of Defense CDMRP PRMRP Discovery Award (PR160710/W81XWH-17-1-0301), and Pilot Grant funding from the School of Health Professions at KU Medical Center (to R.L.M.) and the startup funding in the College of Liberal Arts and Sciences at the University of Kansas (to Y.M.).

## Author Contributions

R.M. and Y.M. designed research; S.P., E. N. M., and B. S. M. performed research; S.P., B. S. M., R.M. and Y.M. analyzed data; and S.P., R.M. and Y.M. wrote the paper.

## References

1. P. C. Harris, V. E. Torres, Genetic mechanisms and signaling pathways in autosomal dominant polycystic kidney disease. The Journal of clinical investigation 124, 2315–2324 (2014).

2. S. M. Nauli et al., Polycystins 1 and 2 mediate mechanosensation in the primary cilium of kidney cells. Nature genetics 33, 129–137 (2003).

3. S. M. Nauli et al., Endothelial cilia are fluid shear sensors that regulate calcium signaling and nitric oxide production through polycystin-1. Circulation 117, 1161–1171 (2008).

4. Z. Pedrozo et al., Polycystin-1 Is a Cardiomyocyte Mechanosensor That Governs L-Type Ca2+ Channel Protein Stability. Circulation 131, 2131–2142 (2015).

5. A. Boletta, G. Germino, Role of polycystins in renal tubulogenesis. Trends Cell Biol 13, 484–492 (2003).

6. C. Piperi, E. K. Basdra, Polycystins and mechanotransduction: From physiology to disease. World journal of experimental medicine 5, 200–205 (2015).

7. Q. Su et al., Structure of the human PKD1-PKD2 complex. Science 361 (2018).

8. Z. Wang et al., The ion channel function of polycystin-1 in the polycystin-1/polycystin-2 complex. EMBO reports 20, e48336 (2019).

9. R. Sandford et al., Comparative analysis of the polycystic kidney disease 1 (PKD1) gene reveals an integral membrane glycoprotein with multiple evolutionary conserved domains. Human molecular genetics 6, 1483–1489 (1997).

10. N. Nims, D. Vassmer, R. L. Maser, Transmembrane domain analysis of polycystin-1, the product of the polycystic kidney disease-1 (PKD1) gene: evidence for 11 membrane-spanning domains. Biochemistry 42, 13035–13048 (2003).

11. S. Kim et al., The polycystin complex mediates Wnt/Ca signalling. Nature cell biology 10.1038/ncb3363 (2016).

12. B. S. Weston, C. Bagneris, R. G. Price, J. L. Stirling, The polycystin-1 C-type lectin domain binds carbohydrate in a calcium-dependent manner, and interacts with extracellular matrix proteins in vitro. Biochimica et biophysica acta 1536, 161–176 (2001).

13. O. Ibraghimov-Beskrovnaya et al., Strong homophilic interactions of the Ig-like domains of polycystin-1, the protein product of an autosomal dominant polycystic kidney disease gene, PKD1. Human molecular genetics 9, 1641–1649 (2000).

14. B. S. Weston, A. N. Malhas, R. G. Price, Structure-function relationships of the extracellular domain of the autosomal dominant polycystic kidney disease-associated protein, polycystin-1. FEBS letters 538, 8–13 (2003).

15. K. Ha et al., The heteromeric PC-1/PC-2 polycystin complex is activated by the PC-1 N-terminus. Elife 9, e60684 (2020).

16. S. C. Parnell et al., The polycystic kidney disease-1 protein, polycystin-1, binds and activates heterotrimeric G-proteins in vitro. Biochemical and biophysical research communications 251, 625–631 (1998).

17. T. Yuasa, A. Takakura, B. M. Denker, B. Venugopal, J. Zhou, Polycystin-1L2 is a novel G-protein-binding protein. Genomics 84, 126–138 (2004).

18. P. Delmas et al., Constitutive activation of G-proteins by polycystin-1 is antagonized by polycystin-2. The Journal of biological chemistry 277, 11276–11283 (2002).

19. S. C. Parnell et al., Polycystin-1 activation of c-Jun N-terminal kinase and AP-1 is mediated by heterotrimeric G proteins. The Journal of biological chemistry 277, 19566–19572 (2002).

20. S. Puri et al., Polycystin-1 activates the calcineurin/NFAT (nuclear factor of activated T-cells) signaling pathway. The Journal of biological chemistry 279, 55455–55464 (2004).

21. P. Delmas et al., Gating of the polycystin ion channel signaling complex in neurons and kidney cells. FASEB journal : official publication of the Federation of American Societies for Experimental Biology 18, 740–742 (2004).

22. Y. Wu et al., Galpha12 is required for renal cystogenesis induced by Pkd1 inactivation. Journal of cell science 129, 3675–3684 (2016).

23. T. Hama, F. Park, Heterotrimeric G protein signaling in polycystic kidney disease. Physiological genomics 48, 429–445 (2016).

24. B. Zhang, U. Tran, O. Wessely, Polycystin 1 loss of function is directly linked to an imbalance in G-protein signaling in the kidney. Development (Cambridge, England) 145 (2018).

25. S. C. Parnell et al., A mutation affecting polycystin-1 mediated heterotrimeric G-protein signaling causes PKD. Human molecular genetics 27, 3313–3324 (2018).

26. C. P. Ponting, K. Hofmann, P. Bork, A latrophilin/CL-1-like GPS domain in polycystin-1. Current biology : CB 9, R585–588 (1999).

27. F. Qian et al., Cleavage of polycystin-1 requires the receptor for egg jelly domain and is disrupted by human autosomal-dominant polycystic kidney disease 1-associated mutations. Proc Natl Acad Sci U S A 99, 16981–16986 (2002).

28. A. Kurbegovic et al., Novel functional complexity of polycystin-1 by GPS cleavage in vivo: role in polycystic kidney disease. Molecular and cellular biology 34, 3341–3353 (2014).

29. R. L. Maser, J. P. Calvet, Adhesion GPCRs as a paradigm for understanding polycystin-1 G protein regulation. Cellular signalling 72, 109637 (2020).

30. D. Arac et al., A novel evolutionarily conserved domain of cell-adhesion GPCRs mediates autoproteolysis. The EMBO journal 31, 1364–1378 (2012).

31. S. Promel, T. Langenhan, D. Arac, Matching structure with function: the GAIN domain of adhesion-GPCR and PKD1-like proteins. Trends in pharmacological sciences 34, 470–478 (2013).

32. Y. Cai et al., Altered trafficking and stability of polycystins underlie polycystic kidney disease. J Clin Invest 124, 5129–5144 (2014).

33. V. G. Gainullin, K. Hopp, C. J. Ward, C. J. Hommerding, P. C. Harris, Polycystin-1 maturation requires polycystin-2 in a dose-dependent manner. The Journal of clinical investigation 125, 607–620 (2015).

34. X. Su et al., Regulation of polycystin-1 ciliary trafficking by motifs at its C-terminus and polycystin-2 but not by cleavage at the GPS site. Journal of cell science 128, 4063–4073 (2015).

35. M. Trudel, Q. Yao, F. Qian, The Role of G-Protein-Coupled Receptor Proteolysis Site Cleavage of Polycystin-1 in Renal Physiology and Polycystic Kidney Disease. Cells 5 (2016).

36. S. Yu et al., Essential role of cleavage of Polycystin-1 at G protein-coupled receptor proteolytic site for kidney tubular structure. Proc Natl Acad Sci U S A 104, 18688–18693 (2007).

37. K. J. Paavola, J. R. Stephenson, S. L. Ritter, S. P. Alter, R. A. Hall, The N terminus of the adhesion G protein-coupled receptor GPR56 controls receptor signaling activity. The Journal of biological chemistry 286, 28914–28921 (2011).

38. K. J. Paavola, R. A. Hall, Adhesion G protein-coupled receptors: signaling, pharmacology, and mechanisms of activation. Molecular pharmacology 82, 777–783 (2012).

39. S. Promel et al., The GPS motif is a molecular switch for bimodal activities of adhesion class G protein-coupled receptors. Cell reports 2, 321–331 (2012).

40. S. C. Petersen et al., The adhesion GPCR GPR126 has distinct, domain-dependent functions in Schwann cell development mediated by interaction with laminin-211. Neuron 85, 755–769 (2015).

41. I. Liebscher et al., A tethered agonist within the ectodomain activates the adhesion G protein-coupled receptors GPR126 and GPR133. Cell reports 9, 2018–2026 (2014).

42. T. Schoneberg, I. Liebscher, R. Luo, K. R. Monk, X. Piao, Tethered agonists: a new mechanism underlying adhesion G protein-coupled receptor activation. Journal of receptor and signal transduction research 35, 220–223 (2015).

43. L. M. Demberg, S. Rothemund, T. Schoneberg, I. Liebscher, Identification of the tethered peptide agonist of the adhesion G protein-coupled receptor GPR64/ADGRG2. Biochemical and biophysical research communications 464, 743–747 (2015).

44. H. M. Stoveken, A. G. Hajduczok, L. Xu, G. G. Tall, Adhesion G protein-coupled receptors are activated by exposure of a cryptic tethered agonist. Proc Natl Acad Sci 112, 6194–6199 (2015).

45. B. S. Magenheimer, E. N. Munoz, J. Ravichandran, R. L. Maser, Constitutive signaling by the C-terminal fragment of polycystin-1 is mediated by a tethered peptide agonist. bioRxiv (2021).

46. Y. Miao, V. A. Feher, J. A. McCammon, Gaussian accelerated molecular dynamics: Unconstrained enhanced sampling and free energy calculation. Journal of chemical theory and computation 11, 3584–3595 (2015).

47. J. Wang et al., Gaussian accelerated molecular dynamics: principles and applications. WIREs Computational Molecular Science 10.1002/wcms.1521, e1521 (2021).

48. A. Roy, A. Kucukural, Y. Zhang, I-TASSER: a unified platform for automated protein structure and function prediction. Nature protocols 5, 725–738 (2010).

49. S. C. Parnell et al., The Polycystic Kidney Disease-1 Protein, Polycystin-1, Binds and Activates Heterotrimeric G-Proteinsin Vitro. Biochemical and biophysical research communications 251, 625–631 (1998).

50. R. H. Purcell, C. Toro, W. A. Gahl, R. A. Hall, A disease-associated mutation in the adhesion GPCR BAI2 (ADGRB2) increases receptor signaling activity. Hum Mutat 38, 1751–1760 (2017).

51. O. Nazarko et al., A Comprehensive Mutagenesis Screen of the Adhesion GPCR Latrophilin-1/ADGRL1. iScience 3, 264–278 (2018).

52. L. Fischer, C. Wilde, T. Schöneberg, I. Liebscher, Functional relevance of naturally occurring mutations in adhesion G protein-coupled receptor ADGRD1 (GPR133). BMC genomics 17, 1–9 (2016).

53. A. Vizurraga, R. Adhikari, J. Yeung, M. Yu, G. G. Tall, Mechanisms of adhesion G protein– coupled receptor activation. Journal of Biological Chemistry 295, 14065–14083 (2020).

54. Y.-Q. Ping et al., Structures of the glucocorticoid-bound adhesion receptor GPR97–G o complex. Nature 589, 620–626 (2021).

55. A. Kishore, R. A. Hall, Disease-associated extracellular loop mutations in the adhesion G protein-coupled receptor G1 (ADGRG1; GPR56) differentially regulate downstream signaling. Journal of Biological Chemistry 292, 9711–9720 (2017).

56. Y. Sun et al., Optimization of a peptide ligand for the adhesion GPCR ADGRG2 provides a potent tool to explore receptor biology. Journal of Biological Chemistry 296, 100174 (2021).

57. X. Han et al., PAR4 activation involves extracellular loop 3 and transmembrane residue Thr153. Blood 136, 2217–2228 (2020).

58. R. L. Maser, B. S. Magenheimer, C. A. Zien, J. P. Calvet, “Transient transfection assays for analysis of signal transduction in renal cells” in Renal Disease. (Springer, 2003), pp. 205–217.

59. Y. Miao, J. A. McCammon, Graded activation and free energy landscapes of a muscarinic G-protein–coupled receptor. Proceedings of the National Academy of Sciences 113, 12162–12167 (2016).

60. J. Huang et al., CHARMM36m: an improved force field for folded and intrinsically disordered proteins. Nature methods 14, 71–73 (2017).

61. J. C. Phillips et al., Scalable molecular dynamics with NAMD. Journal of computational chemistry 26, 1781–1802 (2005).

62. D. A. Case et al., AMBER 18, University of California, San Francisco. (2018).

