## Supporting Information for "Mechanism of Tethered Agonist-Mediated Signaling by Polycystin-1"

**Materials and Methods**

**Materials**

EZ-Link$^{TM}$ Sulfo-NHS-SS-Biotin (21331) and Pierce$^{TM}$ NeutrAvidin$^{TM}$ Agarose were purchased from ThermoFisher Scientific. Dual-Luciferase Reporter Assay Kit was purchased from Promega. Antibodies used for the detection of PC1 include rat IgG1 E8-8C3C10 monoclonal antibody (Baltimore PKD Center; Kerafast), mouse IgG1 7e12 monoclonal and goat polyclonal C20 (Santa Cruz Biotechnology). Rabbit polyclonal antibody against actin (A2066) was purchased from Sigma. Secondary antibodies conjugated to HRP were purchased from Sigma (anti-rabbit, -goat, -mouse) or Jackson ImmunoResearch (anti-rat). Trans-Blot Turbo RTA kit and Clarity$^{TM}$ Western ECL substrate were purchased from BioRad. All other reagents were purchased from Sigma or Fisher.

**CTF expression constructs**

The WT CTF expression construct, consisting of the signal peptide sequence from CD5 joined in frame with PC1 residues 3,049- 4,302, was generated in pCI vector using the full-length human PC1 expression construct AF20 (1), provided by Feng Qian (Addgene #21359). The CD5 signal peptide fragment was generated by PCR from pBS-CD5 (2) using 5'-CD5-Eco and 3'-CD5-BsmBI primers and cloned into pBlueScript (pBS). A fragment starting from the GPS cleavage site and continuing through the stalk and TM1 region of human PC1 (aa residues 3049-3674) was generated by PCR using hCTFXho-BsmBI-For and BsrGIstalk-Rev primers and AF20 as template and then cloned into pBlueScript to generate pBS-HumStalk. The EcoRI-BsmBI fragment of CD5 was subcloned into pBS-

HumStalk to generate pBS-CD5-HumStalk and verified by sequencing. The EcoRI-KpnI fragment of pBS-CD5-HumStalk was then used to replace the EcoRI-KpnI fragment of AF20 (removing residues 1-3,134) to generate the wild-type version of pCI-CD5-HumCTF (WT hCTF). The ΔStalk-CTF construct was generated by the same procedure except for using the HumΔStalk-CTFXho-BsmBI-For primer. Expression constructs with ADPKD-associated missense mutations or simulation substitutions within the stalk region of hCTF (G3052R, R3063C, R3063P, F3066L, N3074A, N3074Q) were made using the QuikChange Lightning Site-Directed Mutagenesis Kit (Agilent Technologies) with pBS-CD5-HumStalk as template, and then used to replace the EcoRI-KpnI fragment of wild-type pCI-CD5-HumCTF. simulation mutants (S3585A, R3835A, R3848E, E4078R) were generated using the mutagenesis kit with pCI-CD5-HumCTF as template, and then used to replace either the TfiI-SphI (S3585A) or the SphI-SapI fragment (R3848E, E4078R) of pCI-CD5-HumCTF. All PCR, mutagenesis and replacement clones were verified by Sanger DNA sequencing (GeneWiz). Primers were synthesized by IDT and sequences are listed in Table S1. In the conduct of research utilizing recombinant DNA, the investigator adhered to NIH Guidelines for research involving recombinant DNA molecules.

**Cell culture, transfection and signaling**

HEK293T cells (ATCC) were maintained and transiently transfected using the Ca-$PO_4$ precipitation method as described previously (3). Briefly, DNA transfection mixes contained NFAT promoter-luciferase reporter (Stratagene), Renilla luciferase (Promega), and either expression vector encoding wild-type or mutant versions of the CTF form of human PC1 (75 ng for signaling or 600 ng for surface biotinylation assays) or an equimolar amount of empty expression vector pCI-neo (ev) as control (37.5 ng or 300 ng). pBlueScript was used to bring the total DNA amount to 8 ug. Cells were lysed in 1X Passive Lysis buffer (PLB; Promega) supplemented with protease and phosphatase inhibitors. Firefly (Fluc)- and Renilla (Rluc)-luciferase activity in each cell lysate was determined using the Dual Luciferase Assay Kit (Promega) and a Berthold tube luminometer. NFAT-Fluc luminescence was normalized to Rluc for each well within a transfection condition, and an average, normalized NFAT-Fluc activity was determined for each condition (n =3 wells/condition). The mean of normalized NFAT-Fluc activity was then used to calculate the Fold NFAT-luc Activation for WT and each CTF mutant relative to the experiment's empty vector control, which were averaged and graphed along with the standard deviation (4) for each expression construct. Transfection experiments were performed a minimum of 3 times (i.e., $\geq$ 3 biological replicates) except where noted.

**Surface biotinylation**

Labeling of cell surface proteins was performed on intact cells with the cell-impermeable, cleavable biotin crosslinking reagent Sulfo-NHS-SS-Biotin at 22-24 hrs following transfection with WT or mutant CTF expression constructs (600 ng). Briefly, cells were rinsed with PBS, and incubated for 30 min at 4˚C in 1.5 mg biotin/ml PBS. Following quenching of the labeling reaction with 50 mM Tris, pH 8.0, cells were scraped off the plate and washed twice with PBS+10 mM Tris before lysis in 1X PLB + protease inhibitors. Cell lysates were spun at 13,000 X g for 3 min. After removing a 10% aliquot of the total cell lysate, biotinylated surface proteins were captured by pulldown with NeutrAvidin agarose beads. Following washes, the beads and final supernatant, (representing cell surface and cytosolic fractions, respectively) were both retained for analysis by Western blot.

**Western blot analyses**

Total and cell surface expression levels of mutant CTF proteins from the signaling and cell surface biotinylation experiments were compared to those of their corresponding WT CTF sample following separation on duplicate, denaturing 7.5% polyacrylamide gels, electrotransfer to nitrocellulose membrane, and probing with primary antibody against PC1 (either C20 or E8; 1:1,000 dilution). Blocking and antibody incubations were performed in Tris-buffered saline-Tween (TBST; 10 mM Tris, 150 mM NaCl, 0.1% Tween 20, pH 7.4) with 5% non-fat dry milk and washes performed with TBST. Detection of HRP-conjugated secondary antibody was performed with Clarity chemiluminescence reagent (BioRad) on an Amersham Imager 600 (GE Healthcare) by capturing multiple exposures. The Amersham ImageQuant TL software was used for quantification of Western blot images. Expression levels of mutant CTF proteins were calculated relative to the WT CTF sample on the same blot, averaged for all experiments and graphed as means with SD error bars.

**Statistical analyses**

All NFAT-luc fold activation and Western blot quantification data are presented as the mean +/- SD from at least 3 independent experiments (except for surface biotinylation analyses of CTF-R3848E and -E4078R expression constructs where n = 2). All statistical comparisons were made in comparison to WT CTF and were performed using a twotailed, unpaired *t* Test calculator from GraphPad (https://www.graphpad.com/quickcalcs/ttest1/?format=SD).

**Gaussian Accelerated Molecular Dynamics (GaMD)**

GaMD is an unconstrained enhanced sampling approach that works by adding a harmonic boost potential to smooth the potential energy surface of biomolecules to reduce energy barriers (5). Brief description of the method is provided here.

Consider a system with $N$ atoms at positions $\vec{r} = \{\vec{r_1}, \ldots, \vec{r_N}\}$. When potential energy of the system $V(\vec{r})$ is less than a threshold energy $E$, a boost potential $\Delta V(\vec{r})$ is added to the system as follows:

$$V^*(\vec{r}) = V(\vec{r}) + \Delta V(\vec{r}), V(\vec{r}) < E \quad (1)$$

$$\Delta V(\vec{r}) = \frac{1}{2}k(E - V(\vec{r}))^2, V(\vec{r}) < E, \quad (2)$$

where $k$ is the harmonic force constant. The two adjustable parameters $E$ and $k$ can be determined by application of three enhanced sampling principles. First, for any two arbitrary potential values $V_1(\vec{r})$ and $V_2(\vec{r})$ found on the original energy surface, if $V_1(\vec{r}) < V_2(\vec{r})$, $\Delta V$ should be a monotonic function that does not change the relative order of the biased potential values, i.e., $V_1^*(\vec{r}) < V_2^*(\vec{r})$. Second, if $V_1(\vec{r}) < V_2(\vec{r})$, the potential difference observed on the smoothed energy surface should be smaller than that of the original, i.e., $V_2^*(\vec{r}) - V_1^*(\vec{r}) < V_2(\vec{r}) - V_1(\vec{r})$. By combining the first two criteria and plugging in the formula of $V^*(\vec{r})$ and $\Delta V$, we obtain:

$$V_{max} \le E \le V_{min} + \frac{1}{k}, \quad (3)$$

where $V_{min}$ and $V_{max}$ are the system minimum and maximum potential energies. To ensure that eq (3) is valid, $k$ has to satisfy: $k \leq \frac{1}{V_{max}-V_{min}}$. Let us define $k \equiv \frac{k_0}{V_{max}-V_{min}}$, then $0 < k_0 \leq 1$. Third, the standard deviation of $\Delta V$ needs to be small enough (i.e., narrow distribution) to ensure accurate reweighting using cumulant expansion to the second order: $\sigma_{\Delta V} = k(E - V_{avg})\sigma_V \leq \sigma_0$, where $V_{avg}$ and $\sigma_V$ are the average and standard deviation of $\Delta V$ with $\sigma_0$ as a user-specified upper limit (e.g., $10k_BT$) for accurate reweighting. When $E$ is set to the lower bound $E = V_{max}$ according to eq 3, $k_0$can be calculated as:

$$k_0 = \min(1.0, k_0') = min\left(1.0, \frac{\sigma_0}{\sigma_V} . \frac{V_{max}-V_{min}}{V_{max}-V_{avg}}\right) \quad (4)$$

Alternatively, when the threshold energy $E$ is set to its upper bound $E = V_{min} + \frac{1}{k}$, $k_0$ is set to:

$$k_0 = k_0^{"} \equiv \left(1 - \frac{\sigma_0}{\sigma_V}\right) . \frac{V_{max}-V_{min}}{V_{avg}-V_{min}} \quad (5)$$

if $k_0^{"}$ is calculated between 0 and 1. Otherwise, $k_0$ is calculated using Eq. (4).

The original GaMD method provides schemes to add only the total potential boost $\Delta V_P$, only dihedral potential boost $\Delta V_D$, or the dual potential boost (both $\Delta V_P$ and $\Delta V_D$). The dual-boost GaMD (GaMD_Dual) simulation generally provides higher acceleration than the other two types of simulations. The simulation parameters comprise of the threshold energy E for applying boost potential and the effective harmonic force constants, $k_{0P}$ and $k_{0D}$ for the total and dihedral potential boost, respectively. Since ligand binding mainly involves nonbonded interactions, another scheme of nonbonded dual-boost GaMD (GaMD_Dual_NB) is introduced in this study of drug binding to the CXCR4 chemokine receptor, for which boost potential is applied to the system dihedral energy and nonbonded potential energy terms.

**Energetic Reweighting of GaMD Simulations**

For energetic reweighting of GaMD simulations to calculate potential mean force (PMF), the probability distribution along a reaction coordinate is written as $p^*(A)$. Given the boost potential $\Delta V(r)$ of each frame, $p^*(A)$ can be reweighted to recover the canonical ensemble distribution $p(A)$, as:

$$p(A_j) = p^*(A_j)\frac{\langle e^{\beta\Delta V(r)}\rangle_j}{\sum_{i=1}^{M}\langle p^*(A_i)e^{\beta\Delta V(r)}\rangle_i}, \quad j = 1,\ldots, M \tag{6}$$

where $M$ is the number of bins , $\beta = k_B T$ and $\langle e^{\beta\Delta V(r)}\rangle_j$ is the ensemble-averaged Boltzmann factor of $\Delta V(r)$ for simulation frames found in the $j^{\text{th}}$ bin. The ensemble-averaged reweighting factor can be approximated using cumulant expansion:

$$\langle e^{\beta\Delta V(r)}\rangle_j = \exp\left\{\sum_{k=1}^{\infty}\frac{\beta^k}{k!}C_k\right\}, \tag{7}$$

where first two cumulants are given by

$$C_1 = \langle\Delta V\rangle,$$

$$C_2 = \langle\Delta V^2\rangle - \langle\Delta V\rangle^2 = \sigma_V^2. \tag{8}$$

The boost potential obtained from GaMD simulations usually follows near-Gaussian distribution. Cumulant expansion to the second order thus provides a good approximation for computing the reweighting factor. The reweighted free energy $F(A) = -k_B T \ln p(A)$ is calculated as

$$F(A) = F^*(A) - \sum_{k=1}^{2} \frac{\beta^k}{k!} C_k + F_c, \quad (9)$$

where $F^*(A) = -k_B T \ln p^*(A)$ is the modified free energy obtained from GaMD simulation and $F_c$ is a constant.

**Homology modeling of PC1 CTF using I-TASSER**

The cryo-EM structure of human PC1-PC2 complex (PDB: 6A70) (6) was used to build the computational model for GaMD simulations. As the protein had several missing regions including the Stalk and several loops, homology modeling of the missing regions was done using I-TASSER web server by providing the sequence of PC1 CTF (chain B) as input. Using template-based modeling approach, two models (model 2 and model 5) out of the top five predicted by I-TASSER gave reasonable conformations of the missing regions with respect to the cryo-EM structure and their orientation in the lipid bilayer. A hybrid model was generated after combining the residues of the two models including model 2 resid 3090-3276 and 3657-3677, and model 5 resid 3049-3089, 3277-3656 and 3678-4168. Resid 3016-3048 of model 5, that consisted of the signal peptide and FLAG epitope tag of the expression vector (6) were deleted along with the large loops connecting TM3-TM4 and TM5-S1 helices as their predicted conformations were not reasonable (**Fig. S1A**). Along with the WT PC1 CTF model, computational models were generated for ΔStalk and the G3052R, R3063C and R3063P ADPKD mutants.

**Simulation system setup**

Computational models were generated using the homology model of PC1 CTF for the WT, ΔStalk by deleting the first 21 residues (3049-3069) of the WT PC1 and the G3052R, R3063C and R3063P ADPKD mutants by single residue mutations in the WT stalk region using PyMOL. The protein was inserted into a palmitoyl-oleoyl-phosphatidyl-choline (POPC) bilayer with all overlapping lipid molecules removed using the *Membrane* plugin in VMD(7). With neutral patches (acetyl and methylamide) added to the termini, the VMD *solvate* plugin (7) was used to neutralize the system in 0.15 M NaCl solution (**Fig. S1B**). GaMD simulations performed on this system are summarized in **Table S2.**

**Simulation protocol**

The CHARMM36m parameter set (8) was used for the PC1 CTF protein and POPC lipid molecules. A similar protocol as used in previous studies (9, 10) was used to prepare the system for GaMD simulations. NAMD2.12 (11) was used for initial energy minimization, followed by thermalization and equilibration. A 12 Å distance cut-off was set for the Van der Waals and short-range electrostatic interactions. Particle-mesh Ewald (12) summation method was used to compute the long-range electrostatic interactions. SHAKE algorithm (13) was implemented on all hydrogen-containing bonds. With all other atoms fixed, the lipid tails were initially energy minimized for 1000 steps using the conjugate gradient algorithm and equilibrated with the constant number, volume and temperature (NVT) ensemble for 0.5 *ns* at 310 K. Further equilibration was performed using the constant number, pressure and temperature (NPT) ensemble at 1 atm and 310 K for 10 *ns* with 5 kcal/(mol·Å$^2$) harmonic position restraints applied to the PDB atoms. Final equilibration of each system was performed using the NPT ensemble at 1 atm and 310 K for 20 *ns* with all atoms unrestrained and a constant ratio constraint applied on the lipid bilayer in the X-Y plane.

The output files of cMD simulations using NAMD were converted to AMBER format using *ParmEd* tool AMBER package (14). GaMD simulations implemented in GPU version of AMBER 18 were performed for the PC1 CTF and its mutant systems. The GaMD simulations involved a short cMD of 12 ns, 48 ns GaMD equilibration followed by 1 μs GaMD production runs for each system using the dual boost GaMD scheme (GaMD_Dual) as listed in **Table S2**.

**Simulation Analysis**

GaMD simulation trajectories were analyzed using CPPTRAJ (15) and VMD (16) tools. Trajectory analysis showed dynamic behavior of the Stalk and the TOP domain of PC1 CTF. A peculiar salt bridge interaction was identified between the TOP domain R3848 and the pore loop E4078 of the PC1 protein. These important interactions were used as reaction coordinates to calculate 2D free energy profiles using the *PyReweighting* toolkit (17). A bin size of 2 Å was used for distances and 10 for the number of contacts. The cutoff of simulation frames in one bin for 2D PMF reweighting (in the range of 500-2000) was set to the minimum number below which the calculated PMF minimum will be shifted.

Residue correlation matrices were calculated from the GaMD simulations as summarized in **Table S2**. During the simulations, each residue with itself will have a correlation value of 1. If two residues moved in the same direction, there was positive correlation between the residues with correlation values in the range of 0 to 1. Alternatively, if two residues moved in the opposite direction, there was negative correlation between the residues with correlation values in the range of -1 to 0.

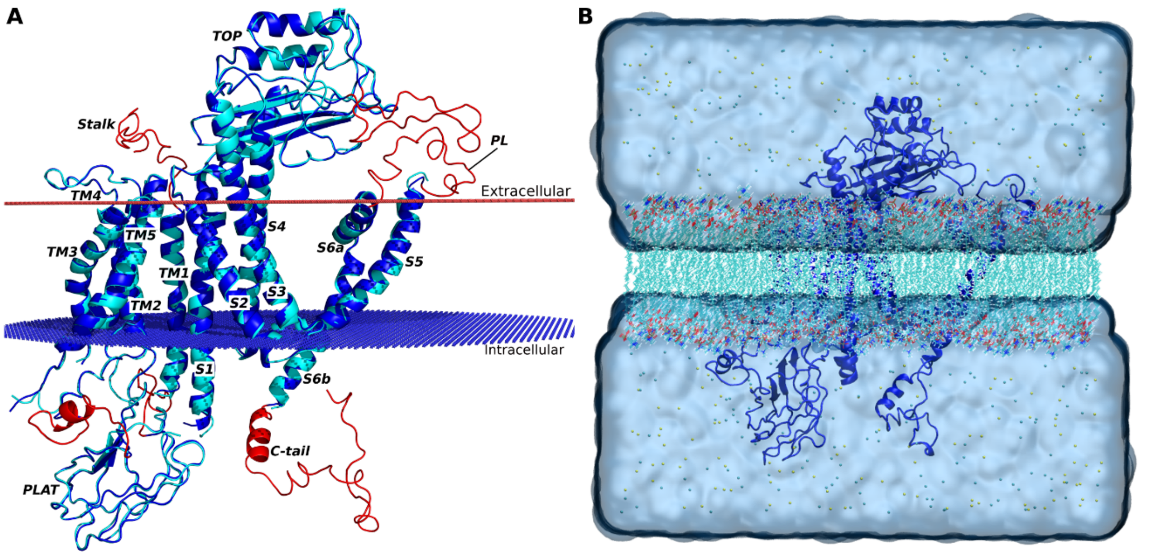


**Figure S1.** (**A**) Computational model of the PC1 CTF extracted from the cryo-EM structure of PC1-PC2 complex (PDB: 6A70), for which the Stalk, Pore Loop (PL) and C-tail missing regions are added through homology modeling. The computational model is represented by blue ribbons with the homology modeled regions highlighted in red. The cryo-EM structure of PC1 CTF is shown in cyan as reference. (**B**) GaMD simulation system of PC1 CTF (blue ribbons) embedded in a POPC lipid bilayer (cyan sticks) solvated in 0.15 M NaCl solution.

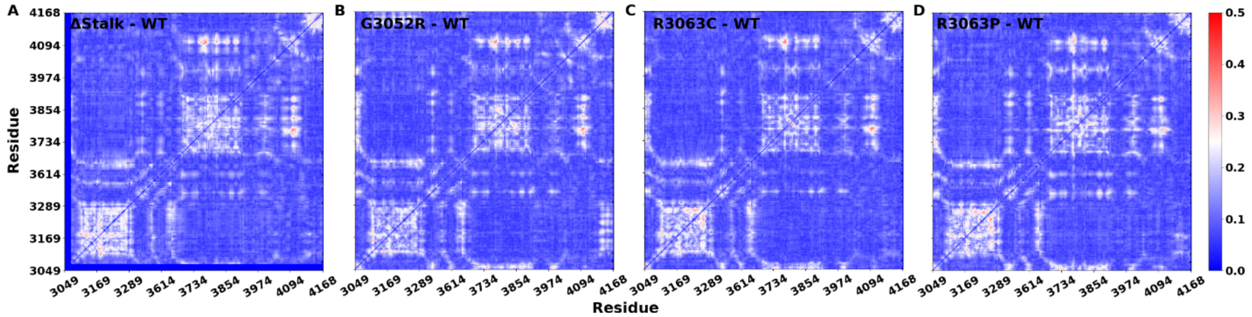


**Figure S2**. The standard errors of differences in residue correlations between the mutant and WT CTF systems calculated from the GaMD simulations: (A) ΔStalk, (B) G3052R, (C) R3063C and (D) R3063P systems. The corresponding average differences in residue correlations between the mutant and WT CTF systems were plotted in **Figs. 2B-2D** and **4A**.

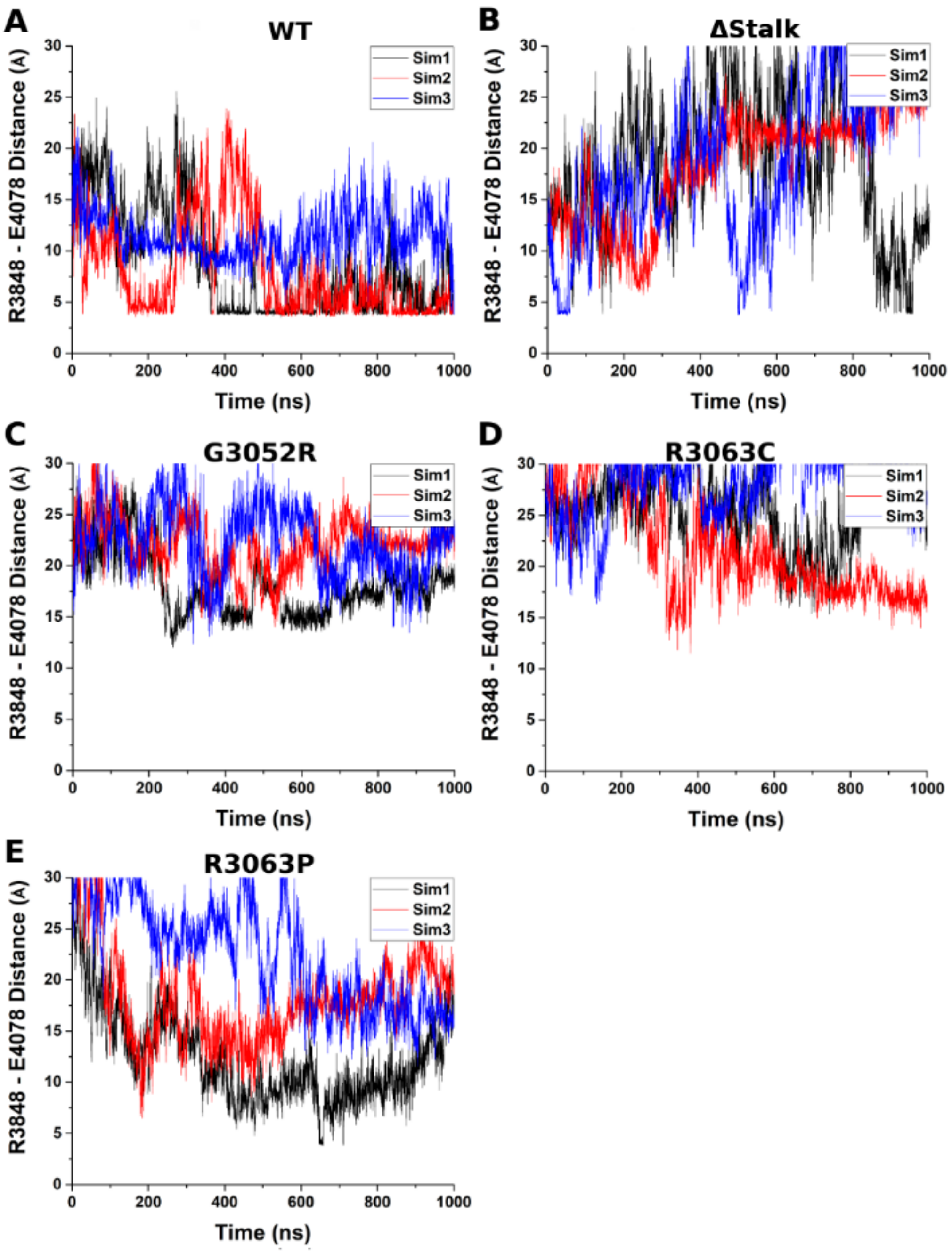


**Figure S3**. Time courses of the distance between the charge centers of the TOP domain residue R3848 (the CZ atom) and PL residue E4078 (the CD atom) calculated from GaMD simulations of the (**A**) WT, (**B**) ΔStalk, (**C**) G3052R, (**D**) R3063C, and (**E**) R3063P PC1 CTF systems.

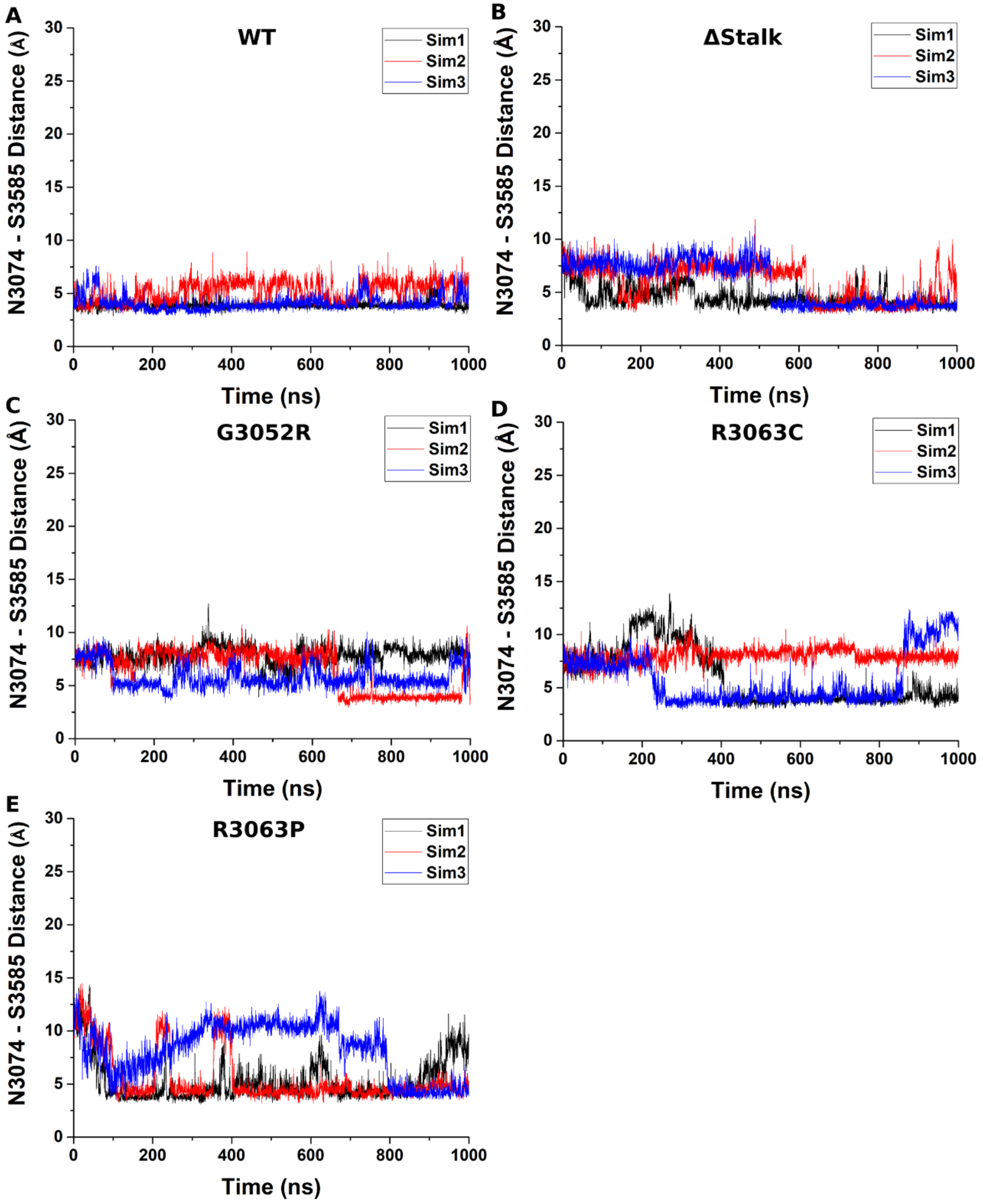


**Figure S4.** Time courses of the distance between the Stalk residue N3074 (the CG atom) and TM5 residue S3585 (the OG atom) calculated from GaMD simulations of the (**A**) WT, (**B**) ΔStalk, (**C**) G3052R, (**D**) R3063C, and (**E**) R3063P PC1 CTF systems.

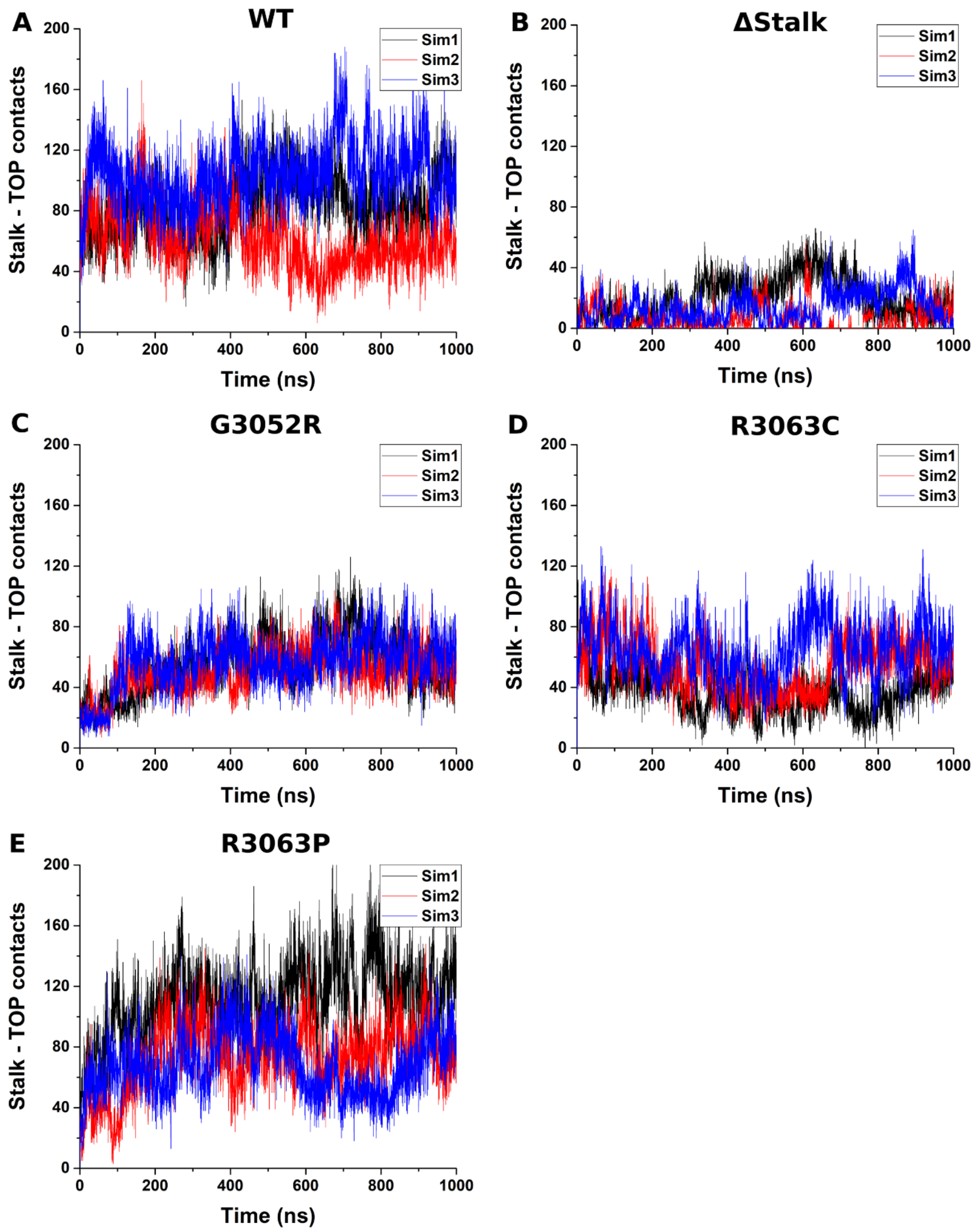


**Figure S5.** Time courses of the number of contacts between the Stalk and TOP domains calculated from GaMD simulations of the (**A**) WT, (**B**) ΔStalk, (**C**) G3052R, (**D**) R3063C, and (**E**) R3063P PC1 CTF systems.

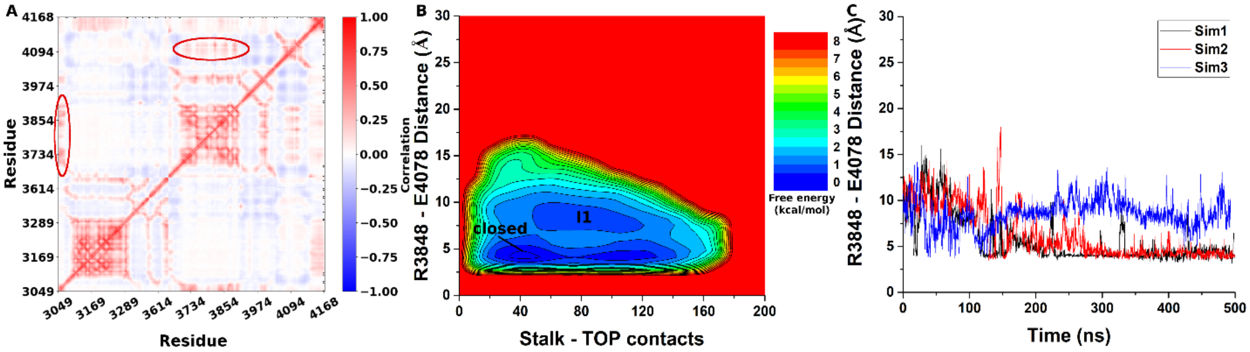


**Figure S6**. (**A**) The average correlation matrix of residue motions calculated from three independent GaMD simulations using the AMBER FF19SB force field of the WT PC1 CTF. High correlations between Stalk-TOP and TOP-PL domains are highlighted in red circles. (**B**) 2D free energy profiles regarding the number of atom contacts between the Stalk and TOP domains and the R3848-E4078 distance (the CZ atom in R3848 and the CD atom in E4078) calculated from the GaMD simulations. Residue contacts were calculated for heavy atoms with a distance cutoff of 4 Å. Important low-energy conformational states are identified, including the "Closed" and "Intermediate I1". (**C**) Time course of the distance between the charge centers of the TOP domain residue R3848 (the CZ atom) and PL residue E4078 (the CD atom) calculated from GaMD simulations.

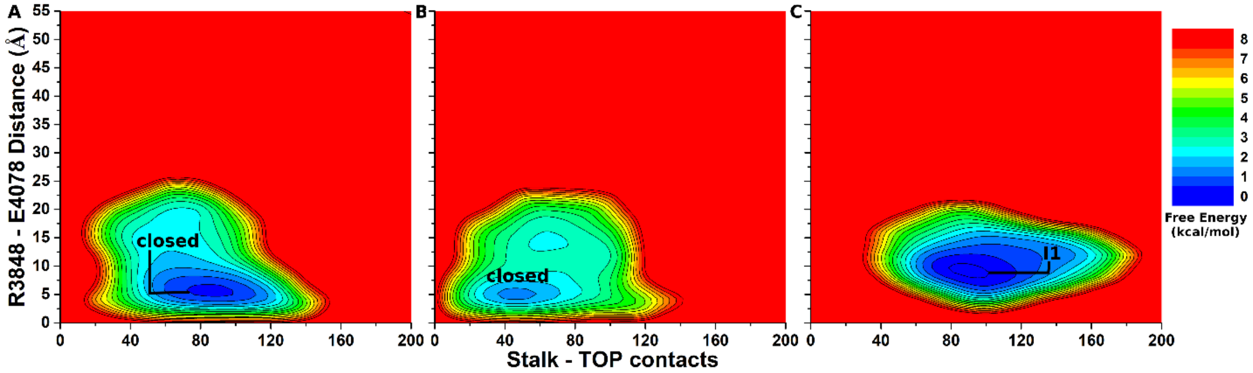


**Figure S7.** 2D free energy profiles of the WT PC1 CTF system regarding the number of atom contacts between the Stalk and TOP domains and the R3848-E4078 distance (the CZ atom in R3848 and the CD atom in E4078) calculated from (**A**) Sim1, (**B**) Sim2, and (**C**) Sim3 of the GaMD simulations. Important low-energy conformational states are identified, including the "Closed" and "Intermediate I1".

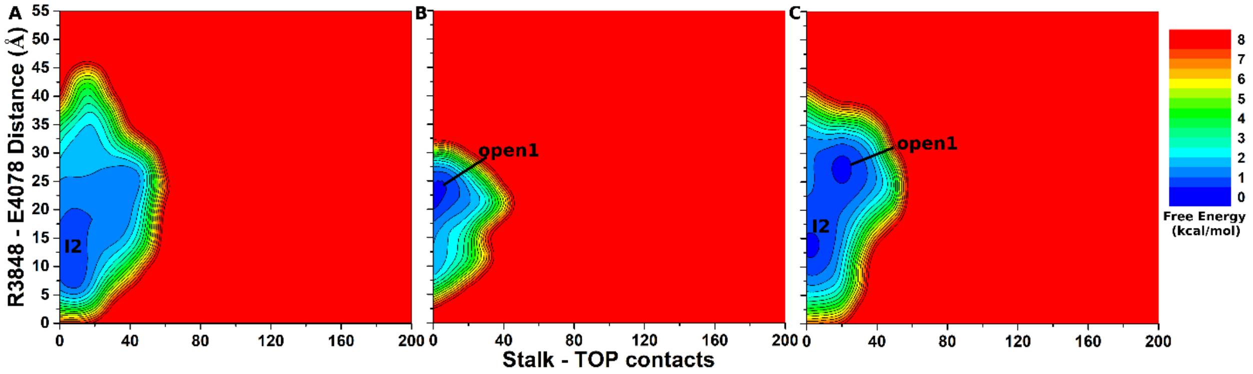


**Figure S8.** 2D free energy profiles of the ΔStalk PC1 CTF system regarding the number of atom contacts between the Stalk and TOP domains and the R3848-E4078 distance (the CZ atom in R3848 and the CD atom in E4078) calculated from (**A**) Sim1, (**B**) Sim2, and (**C**) Sim3 of the GaMD simulations. Important low-energy conformational states are identified, including the "Open1" and "Intermediate I2".

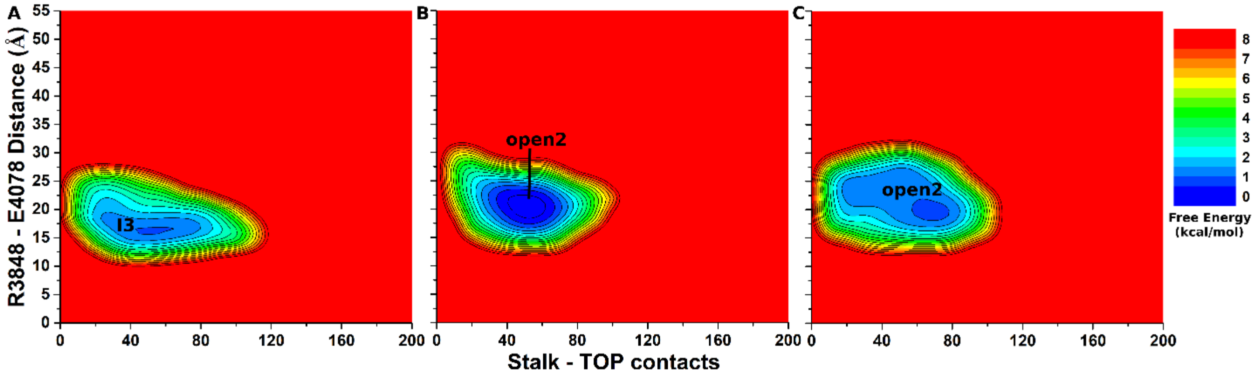


**Figure S9.** 2D free energy profiles of the G3052R PC1 CTF system regarding the number of atom contacts between the Stalk and TOP domains and the R3848-E4078 distance (the CZ atom in R3848 and the CD atom in E4078) calculated from (**A**) Sim1, (**B**) Sim2, and

(**C**) Sim3 of the GaMD simulations. Important low-energy conformational states are identified, including the "Open2" and "Intermediate I3".

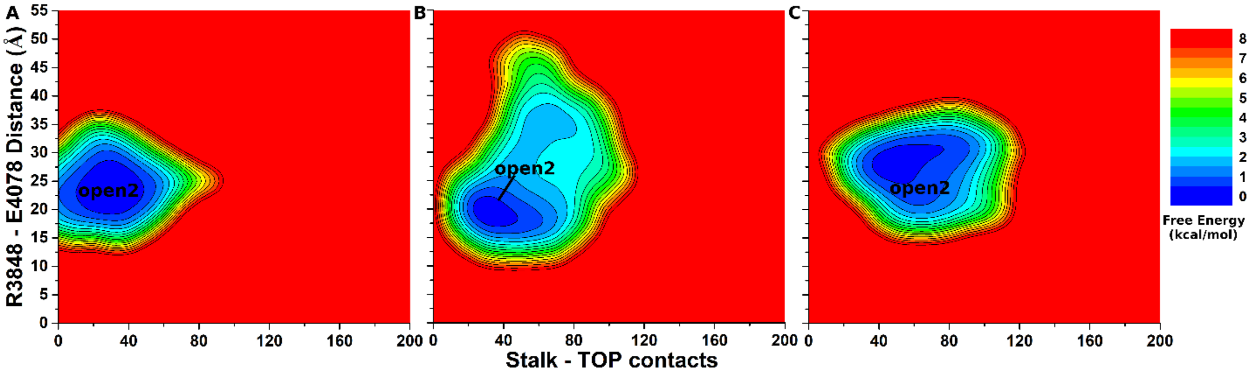


**Figure S10.** 2D free energy profiles of the R3063C PC1 CTF system regarding the number of atom contacts between the Stalk and TOP domains and the R3848-E4078 distance (the CZ atom in R3848 and the CD atom in E4078) calculated from (**A**) Sim1, (**B**) Sim2, and (**C**) Sim3 of the GaMD simulations. "Open2" low-energy conformational states are identified.

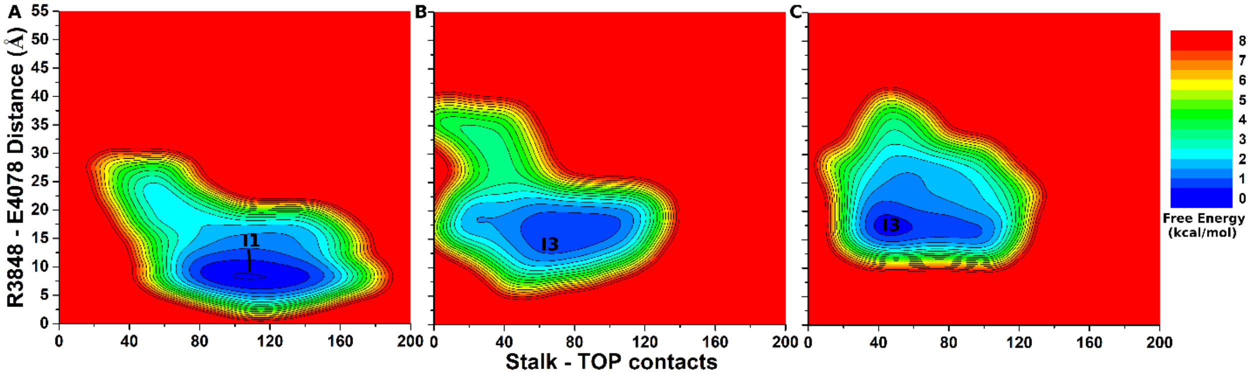


**Figure S11.** 2D free energy profiles of the R3063P PC1 CTF system regarding the number of atom contacts between the Stalk and TOP domains and the R3848-E4078 distance (the CZ atom in R3848 and the CD atom in E4078) calculated from (**A**) Sim1, (**B**) Sim2, and (**C**) Sim3 of the GaMD simulations. Important low-energy conformational states are identified, including the "Intermediate I1" and "Intermediate I3".

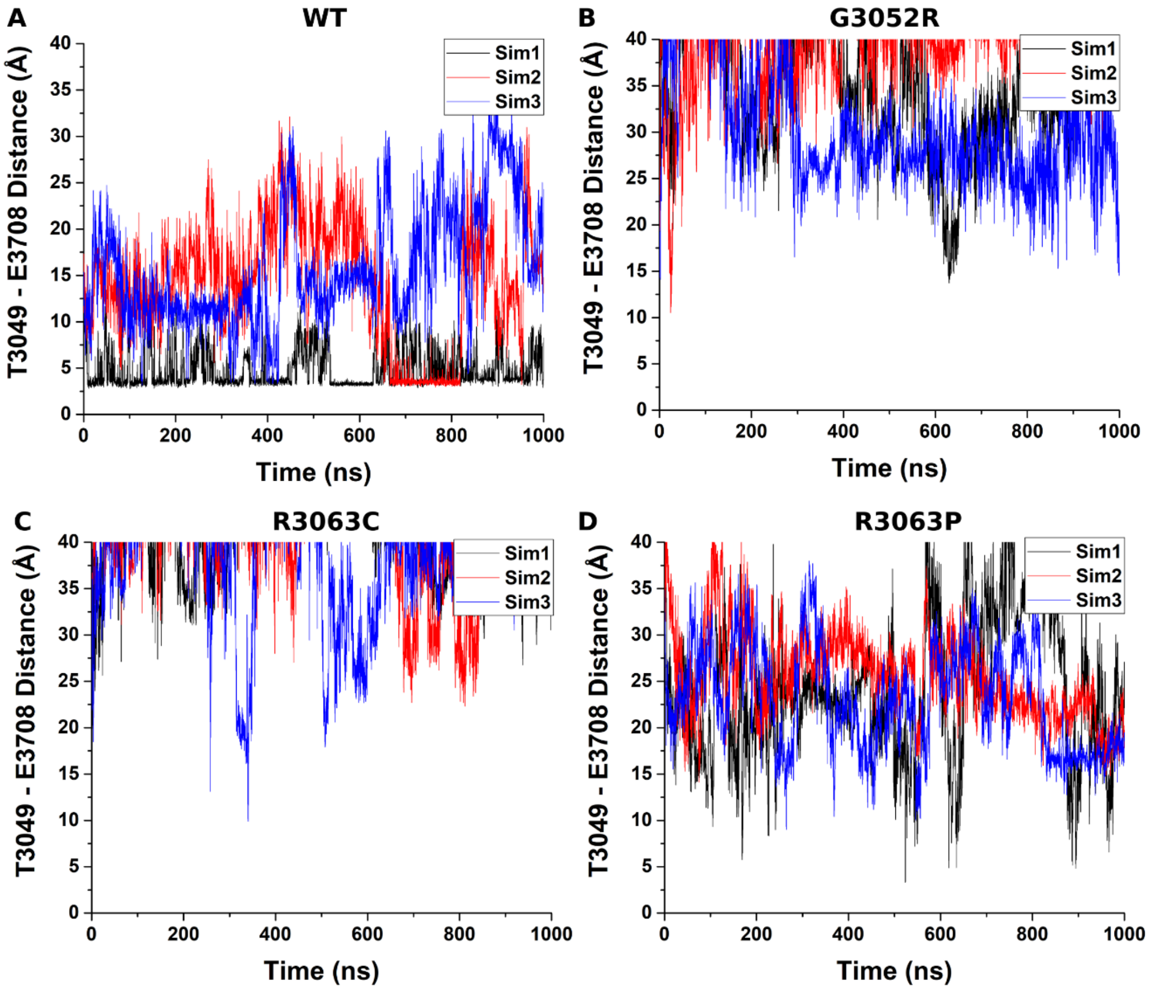


**Fig. S12.** Time courses of the distance between the Stalk residue T3049 (the OG1 atom) and TOP residue E3078 (the CD atom) calculated from GaMD simulations of the (**A**) WT, (**B**) G3052R, (**C**) R3063C, and (**D**) R3063P PC1 CTF systems.

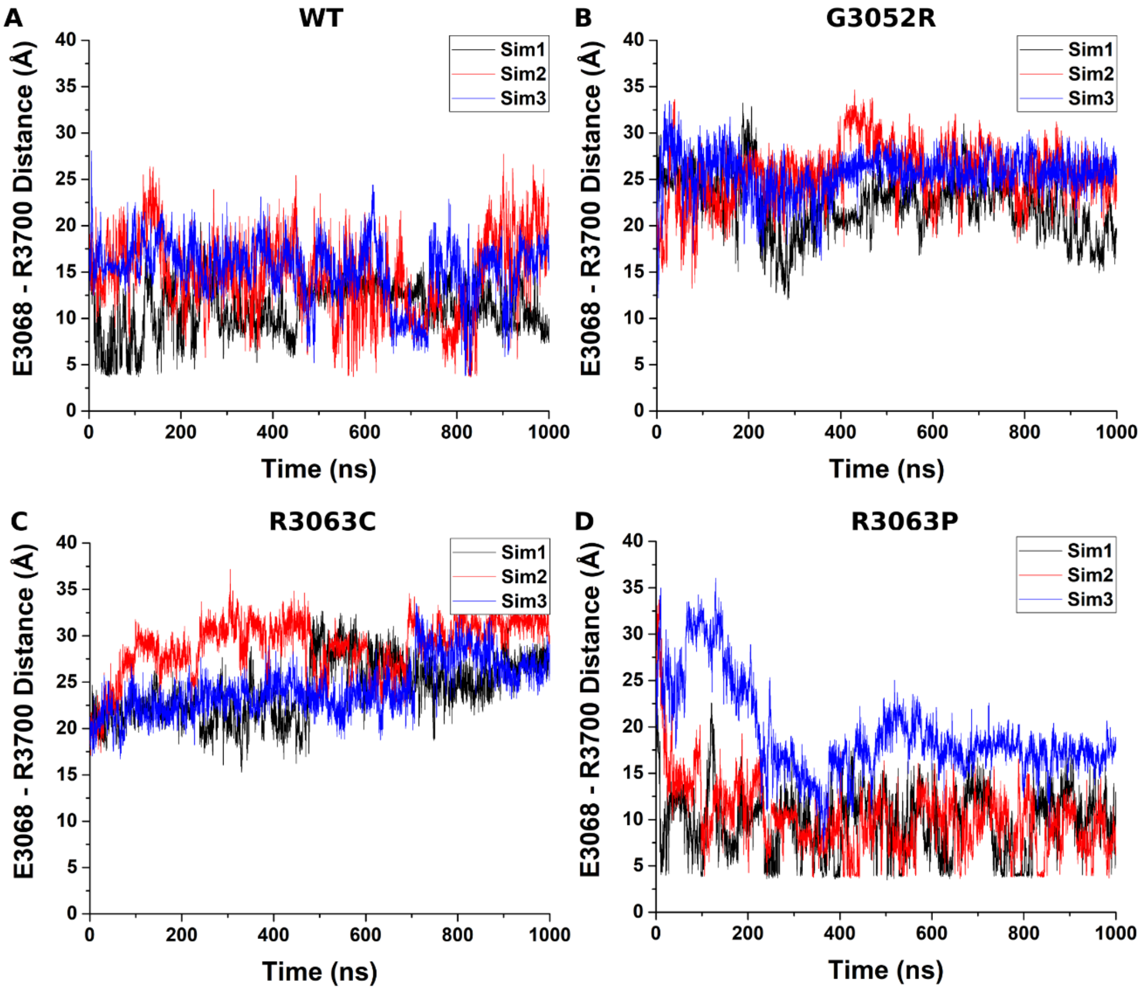


**Fig. S13.** Time courses of the distance between the charge centers of the Stalk residue E3068 (the CD atom) and TOP residue R3700 (the CZ atom) calculated from GaMD simulations of the (**A**) WT, (**B**) G3052R, (**C**) R3063C, and (**D**) R3063P PC1 CTF systems.

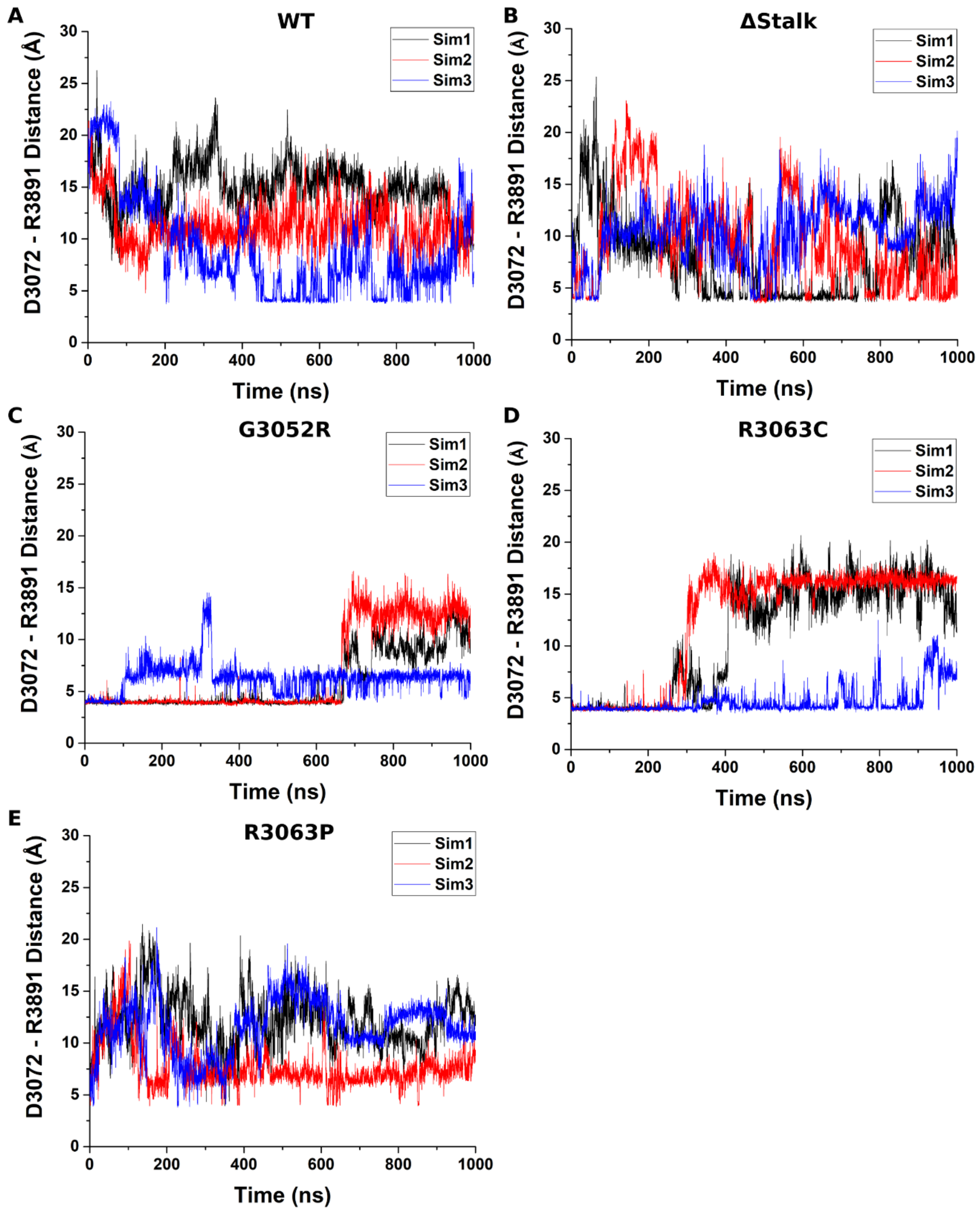


**Fig. S14.** Time courses of the distance between the charge centers of the Stalk residue D3072 (the CG atom) and TOP residue R3891 (the CZ atom) calculated from GaMD simulations of the (**A**) WT, (**B**) ΔStalk, (**C**) G3052R, (**D**) R3063C, and (**E**) R3063P PC1 CTF systems.

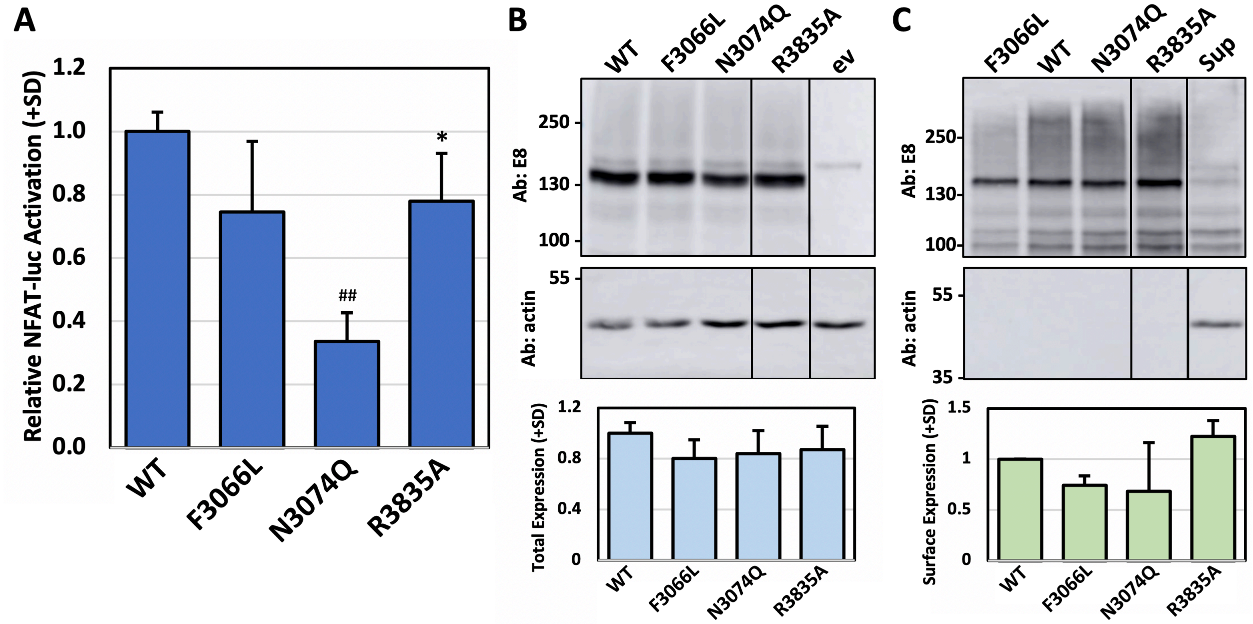


**Figure S15. Effect of proposed 'neutral mutations' on CTF-mediated NFAT reporter activation.** (A) Average relative NFAT reporter activation levels (+SD) of neutral CTF mutants, F3066L, N3074Q, and R3835A, normalized to WT CTF from 2-4 separate transfection experiments with n=3 wells/expression construct/experiment. *, $p<0.05$; ##, $p<0.0001$ relative to WT CTF. (B) Representative Western blot of the WT and neutral CTF mutants from one of the experiments represented in (A). Image is from a single blot. Lines between lanes indicate removal of a non-relevant, intervening lane. Blot was first probed with E8 antibody to detect CTF proteins (top blot) and then reprobed with an antibody against actin (bottom blot). Graph below represents the average total expression level (+SD) for the neutral mutant CTF proteins relative to WT CTF from the experiments in (A). (C) Representative Western blot of the surface biotinylation analyses of the WT and neutral mutant CTF proteins from two of the experiments in (A). Supernatant (Sup) lane is from the pulldown with WT CTF. Image is from a separate blot. Lines between lanes indicate removal of a non-relevant, intervening lane. Blot was first probed with E8 antibody to detect CTF proteins (top blot) and then reprobed with an antibody against actin (bottom blot) to show cleanliness of surface protein biotinylation. Graph below shows the average relative cell surface expression level (+SD) for WT and neutral mutant CTF proteins from 2 separate surface biotinylation experiments.

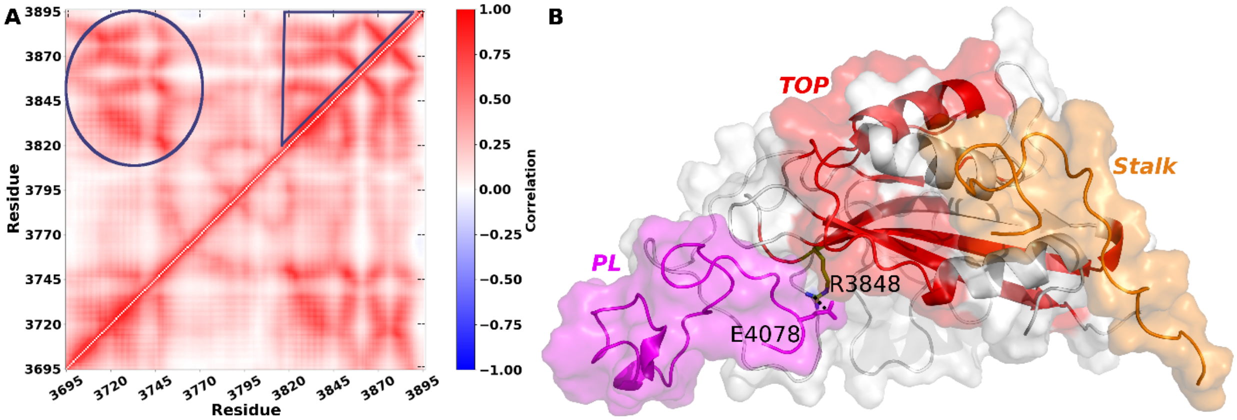


**Figure S16.** (**A**) The average correlation matrix of residue motions in the TOP domain calculated from three independent GaMD simulations of the WT PC1 CTF. High correlations within the TOP domain are highlighted with the blue circle and triangle. (**B**) Cartoon representation of the potential signal propagation between the Stalk-TOP and TOP-PL domains of the WT PC1 CTF. Structural elements within the TOP domain with increased correlation of residue motions are shown in red. The Stalk (orange), TOP domain (white) and pore loop (magenta) are labeled in the PC1 CTF. The R3848-E4078 residue interaction is represented in sticks.

**Table S1. Primer Sequences for PCR Cloning and Mutagenesis.**

| Primer | Sequence |
|---|---|
| 5’-CD5 Eco | 5’- TTCTAGAATTCCCTCGACCTCG -3’ |
| 3’-CD5-BsmBI | 5’- GACTAGCGTCTCATGCCTAGCACGGAAGC -3’ |
| hCTF Xho-BsmBI For | 5’-GACTAGCTCGAGCGTCTCAGGCACCGCCTTCGGCGCCAGCCTCTTC-3’ |
| BsrGIstalk-Rev | 5’- AGGGTCTGGGTAGAGTGCTT -3’ |
| HumΔStalk-CTF Xho-BsmBI For | 5’-GACTAGCTCGAGCGTCTCAGGCACAGCGGATGTAAACTAC-3’ |
| G3052R Top | 5’-CACCGCCTTCCGCGCCAGCCTC-3’ |
| G3052R Bott | 5’-GAGGCTGGCGCGGAAGGCGGTG-3’ |
| R3063C For | 5’-GCCCCCAAGCCATGTCTGCTTTGTGTTTCCTGAG-3’ |
| R3063C Rev | 5’-CTCAGGAAACACAAAGCAGACATGGCTTGGGGGC-3’ |
| R3063P Top | 5’-GCCCCCAAGCCATGTCCCCTTTGTGTTTCCTGAG-3’ |
| F3066L Top | 5’-GTCCGCTTTGTGTTACCTGAGCCGACAGC-3’ |
| F3066L Bott | 5’- GCT GTC GGC TCA GGT AAC ACA AAG CGG AC-3’ |
| R3063P Bott | 5’-CTCAGGAAACACAAAGGGGACATGGCTTGGGGGC-3’ |
| N3074A For | 5’-GAGCCGACAGCGGAT GTA GCCTACATCGTCATGCTGAC-3’ |
| N3074A Rev | 5’-GTCAGCATGACGATGTAGGCTACATCCGCTGTCGGCTC-3’ |
| N3074Q Top | 5’- GAGCCGACAGCGGATGTACAATACATCGTCATGCTGAC-3’ |
| N3074Q Bott | 5’- GTCAGCATGACGATGTATTGTACATCCGCTGTCGGCTC-3’ |
| S3585A For | 5’-CCCCCCGGGCGTGGCTGTTGCGTGGCTCC-3’ |
| S3585A Rev | 5’-GGAGCCACGCAACAGCCACGCCCGGGGGG-3’ |
| R3835A Top | 5’- CCGCGACCGGCTGGCCTTCCTGCAGCTG-3’ |
| R3835A Bott | 5’- CAGCTGCAGGAAGGCCAGCCGGTCGCGG-3’ |
| R3848E Top | 5’-TGGCTGGACAACAGGAGCGAGGCTGTGTTCCTGGAGCTC-3’ |
| R3848E Bott | 5’-GAGCTCCAGGAACACAGCCTCGCTCCTGTTGTCCAGCCA-3’ |
| E4078R Top | 5’-CCCTGTGTCCTGCCAGGTCCTGGCACCTGTC-3’ |
| E4078R Bott | 5’-GACAGGTGCCAGGACCTGGCAGGACACAGGG-3’ |

**Table S2. Summary of GaMD simulations performed on the WT, ΔStalk and the G3052R, R3063C and R3063P ADPKD mutants of PC1 CTF.**

| System | $N_{atoms}$ [a] | ID | Length (ns) | $\Delta V_{avg}$ [b] (kcal/mol) | $\sigma_{\Delta V}$ [c] (kcal/mol) |
|---|---|---|---|---|---|
| **WT** | 269,877 | Sim1 | 1000 | 24.13 | 7.25 |
| | | Sim2 | | 24.12 | 7.25 |
| | | Sim3 | | 24.37 | 7.27 |
| **ΔStalk** | 269,887 | Sim4 | | 22.78 | 7.20 |
| | | Sim5 | | 23.17 | 7.23 |
| | | Sim6 | | 23.05 | 7.23 |
| **G3052R** | 269,880 | Sim7 | | 24.95 | 7.91 |
| | | Sim8 | | 25.16 | 7.93 |
| | | Sim9 | | 24.43 | 7.84 |
| **R3063C** | 269,875 | Sim10 | | 22.93 | 7.17 |
| | | Sim11 | | 22.91 | 7.17 |
| | | Sim12 | | 23.02 | 7.18 |
| **R3063P** | 269,878 | Sim13 | | 25.44 | 6.67 |
| | | Sim14 | | 25.33 | 6.65 |
| | | Sim15 | | 25.36 | 6.66 |

**[a] $N_{atoms}$: number of atoms in the system**

**[b] $\Delta V_{avg}$: average of the GaMD boost potential**

**[c] $\sigma_{\Delta V}$: standard deviation of the GaMD boost potential**

**References:**


1. F. Qian *et al.*, Cleavage of polycystin-1 requires the receptor for egg jelly domain and is disrupted by human autosomal-dominant polycystic kidney disease 1-associated mutations. *Proceedings of the National Academy of Sciences of the United States of America* **99**, 16981-16986 (2002).
2. N. Nims, D. Vassmer, R. L. Maser, Transmembrane domain analysis of polycystin-1, the product of the polycystic kidney disease-1 (PKD1) gene: evidence for 11 membrane-spanning domains. *Biochemistry* **42**, 13035-13048 (2003).
3. R. L. Maser, B. S. Magenheimer, C. A. Zien, J. P. Calvet, "Transient transfection assays for analysis of signal transduction in renal cells" in Renal Disease. (Springer, 2003), pp. 205-217.
4. G. Dalagiorgou *et al.*, Mechanical stimulation of polycystin-1 induces human osteoblastic gene expression via potentiation of the calcineurin/NFAT signaling axis. *Cell Mol Life Sci* **70**, 167-180 (2013).
5. Y. Miao, V. A. Feher, J. A. McCammon, Gaussian accelerated molecular dynamics: Unconstrained enhanced sampling and free energy calculation. *Journal of chemical theory and computation* **11**, 3584-3595 (2015).
6. Q. Su *et al.*, Structure of the human PKD1-PKD2 complex. *Science* **361** (2018).
7. W. Humphrey, A. Dalke, K. Schulten, VMD: Visual molecular dynamics. *Journal of Molecular Graphics & Modelling* **14**, 33-38 (1996).
8. J. Huang *et al.*, CHARMM36m: an improved force field for folded and intrinsically disordered proteins. *Nature methods* **14**, 71-73 (2017).
9. Y. Miao, J. A. McCammon, Graded activation and free energy landscapes of a muscarinic G-protein–coupled receptor. *Proceedings of the National Academy of Sciences* **113**, 12162-12167 (2016).
10. Y. Miao, J. A. McCammon, Mechanism of the G-protein mimetic nanobody binding to a muscarinic G-protein-coupled receptor. *Proceedings of the National Academy of Sciences* **115**, 3036-3041 (2018).
11. J. C. Phillips *et al.*, Scalable molecular dynamics with NAMD. *Journal of computational chemistry* **26**, 1781-1802 (2005).
12. T. Darden, D. York, L. Pedersen, Particle mesh Ewald: An N· log (N) method for Ewald sums in large systems. *The Journal of chemical physics* **98**, 10089-10092 (1993).
13. J.-P. Ryckaert, G. Ciccotti, H. J. Berendsen, Numerical integration of the cartesian equations of motion of a system with constraints: molecular dynamics of n-alkanes. *Journal of computational physics* **23**, 327-341 (1977).
14. D. A. Case *et al.*, AMBER 18, University of California, San Francisco. (2018).
15. D. R. Roe, T. E. Cheatham III, PTRAJ and CPPTRAJ: software for processing and analysis of molecular dynamics trajectory data. *J Chem Theory Comput* **9**, 3084-3095 (2013).
16. W. Humphrey, A. Dalke, K. Schulten, VMD: visual molecular dynamics. *Journal of molecular graphics* **14**, 33-38 (1996).
17. Y. Miao *et al.*, Improved reweighting of accelerated molecular dynamics simulations for free energy calculation. *Journal of Chemical Theory and Computation* **10**, 2677-2689 (2014).
